# M-GWAS for the gut microbiome in Chinese adults illuminates on complex diseases

**DOI:** 10.1101/736413

**Authors:** Xiaomin Liu, Shanmei Tang, Huanzi Zhong, Xin Tong, Zhuye Jie, Qiuxia Ding, Dan Wang, Ruidong Guo, Liang Xiao, Xun Xu, Huanming Yang, Jian Wang, Yang Zong, Xiao Liu, Yong Zhang, Susanne Brix, Karsten Kristiansen, Yong Hou, Huijue Jia, Tao Zhang

## Abstract

The gut microbiome has been established as a key environmental factor to health. Genetic influences on the gut microbiome have been reported, yet, doubts remain as to the significance of genetic associations. Here, we provide shotgun data for whole genome and whole metagenome from a Chinese cohort, identifying no less than 20% genetic contribution to the gut microbiota. Using common variants-, rare variants- and copy number variations (CNVs)-based association analyses, we identified abundant signals associated with the gut microbiome especially in metabolic, neurological and immunological functions. The controversial concept of enterotypes may have a genetic attribute, with the top 2 loci explaining 11% of the *Prevotella-Bacteroides* variances. Stratification according to gender led to the identification of differential associations in males and females. Genetically encoded responses to ectopic presence of oral bacteria in the gut appear to be a common theme in a number of diseases investigated by MWAS (Metagenome-wide association studies). Our two-stage M-GWAS (Microbiome genome-wide association studies) on a total of 1295 individuals unequivocally illustrates that neither microbiome nor GWAS studies could overlook one another in our quest for a better understanding of human health and diseases.

**Highlights:** M-GWAS using high-depth whole genome identifies contributions from rare variants and CNVs.

Gut microbial modules such as butyrate, amino acids, mucin degradation show genetic associations.

Gender differential M-GWAS underscores differences in metabolic and psychological predispositions.

Some of the MWAS markers for colorectal cancer and cardiometabolic diseases show genetic associations.

## Introduction

The gut microbiome is increasingly recognized as playing important roles in host health and diseases, exerting influences well beyond the gut (Blacher et al., 2019; Wang and Jia, 2016). However, due to modulations by diet and medication, the gut microbiome is commonly viewed as highly dynamic, whereas disease markers are expected to be stable. Studies in mice (Org et al., 2015) and in human twins (Goodrich et al., 2016; Xie et al., 2016) have observed substantial heritability for some bacteria. Several genome-wide association analysis studies (Blekhman et al., 2015; Bonder et al., 2016; Rothschild et al., 2018; Turpin et al., 2016; Wang et al., 2016) for the gut microbiota (M-GWAS) have found associations between host single nucleotide polymorphisms (SNPs) and individual bacterial taxa, beta diversity or pathways. Yet, doubts remain as to the significance of genetic associations. For example, a recent study (Rothschild et al., 2018) including a heterogeneous population of ∼800 individuals reported that the average heritability of gut microbiome taxa is only 1.9%, in contrast to Wang et al. which identified 42 SNPs that together explained 10% of the variance of the β-diversity (Wang et al., 2016). In addition, these studies all used genotyping arrays data for host genome and they all used 16S rRNA gene amplicon sequencing except for one study which had low-depth shotgun data for fecal samples. The lack of high-depth whole genome sequencing (WGS) data means that the studies rely on imputation for SNPs and could be missing potential associations from insertions/deletions (indels), copy number variations (CNVs), especially for rare variants. In addition, previous GWAS studies for the gut microbiome investigated populations of European ancestry, how host genetics shape the gut microbiome in an Asian population remains unknown.

In this study, we identified genetic-microbial associations using for the first time high-depth sequencing data for both whole-genome and metagenome, in a high-depth discovery cohort of 632 healthy Chinese individuals and a low-depth validation cohort of 663 individuals. *Prevotella* and *Bifidobacterium* are the most heritable taxa after the cellulose-degrading phylum Fibrobacteres in this Chinese cohort, in agreement with twins results from European (Xie et al., 2016) as well as Asian (Lim et al., 2017) cohorts. With WGS data, we are uniquely positioned to comprehensively investigate common variants, rare variants, CNVs and the HLA (human leukocyte antigen) locus associated with the gut microbiome. Considering the reported gender differences in the gut microbiome (Xiao et al., 2016; Xiao et al., 2015) and increasing interest in incorporating the gender perspective into metagenomic and genomic studies (Khramtsova et al., 2018), we carry out the first gender-specific M-GWAS to understand the difference of gut microbiome-genome association between genders. Together, our results reveal host genetic influences on the composition and functional potential of the gut microbiome to an unprecedented extent and generate a number of testable hypothesis for diseases such as colorectal cancer and cardiometabolic diseases.

## Results

### Characteristics not reported in European cohorts

To investigate genetic influences on the gut microbiome, we performed whole-genome sequencing on 632 blood samples to a mean depth of 44× (range from 32× to 52×, **Fig. S1A, Table S1A**) per individual, and metagenomic sequencing on 632 stool samples to an average of 8.57 ± 2.21 GB (**Fig. S1B**). This discovery cohort had a mean age of 30.7 ± 5.5 years (mean ± s.d.; range of 6–35 years), a mean body mass index (BMI) of (21.8 ± 6.3) and 53.5% were female (**Table S1B**). We observed in this Chinese cohort that each genome differs between one another by 3.9 to 4.9 million sites (**Table S1C**). Variants were directly determined from the high-depth human genomes, including 38 million SNPs, 5 million indels, and 40 thousand CNVs. 6.5 million of these were common variants (minor allele frequency (MAF) > 0.05), 36.5 million of these were rare and low-frequency variants (MAF) ≤ 0.05). Taxonomic profiling of the fecal metagenomes resulted in 19 phyla, 21 classes, 40 order, 77 families, 307 genera and 519 species. The top five abundant phyla in this cohort were Bacteroidetes (relative abundance of 51.0 ± 13.5%), Firmicutes (11.2 ± 5.6%), Proteobacteria (2.8 ± 3.7%), Fusobacteria (0.3 ± 1.1%) and Actinobacteria (0.13 ± 0.27%) (**Fig. S2**). Based on existing knowledge, we performed all M-GWAS by including covariates for gender, age, BMI, diet and lifestyle factors, stool form, defecation frequency, as well as the top four principal components to account for population structure (**Table S1B; methods**).

Unlike M-GWAS using chip data from European cohorts, we identified suggestive host genetic associations for the controversial concept of enterotypes (Fig. 1, **Table S1D**), possibly due to higher prevalence of *Prevotella* in developing countries (Dhakan et al., 2019; Vangay et al., 2018). Principle coordinate analysis (PCoA) as well as Dirichlet multinominal mixture model using Bray-Curtis dissimilarity showed that this Chinese cohort could be formed into two clusters dominated by *Bacteroides* and *Prevotella* (Zou et al., 2019), containing 440 and 178 individuals, respectively (Fig. 1A). The top two loci associated with the *Bacteroides*-*Prevotella* dichotomy (*P_P-B_* = 2.08 × 10^−6^ and *P_P-B_* = 2.6 × 10^−6^, respectively, using *Prevotella* as cases and *Bacteroides* as controls in a logistic regression model) together explained 11% of the variance of the *Bacteroides* versus *Prevotella* ‘enterotype’. Despite a report challenging the negative association between *Bacteroides* and *Prevotella* (Vandeputte et al., 2017), due to the statistically well-known loss of one degree of freedom in compositional data (Cao et al., 2019), genetic associations for these genera also showed the opposite trend. The minor allele of the top SNP, rs13045408 at *BTBD3*-*LINC01722*, positively correlated with *Bacteroides* abundance (β = 0.043; P = 5.3 × 10^−3^) and negatively correlated with *Prevotella* abundance (β = −1.76; P = 1.6 × 10^−4^) (*P_P-B_* = 2.1 × 10^−6^; Fig. 1C); on the other hand, the minor allele of the other SNP rs1453213 at *OXR1* positively correlated with *Prevotella* (β= 2.23; P =1.3 × 10^−7^) and negatively correlated with *Bacteroides* (β = − 0.049; P = 3.4 × 10^−4^) (*P_P-B_* = 2.6 × 10^−6^).

**Figure 1.**
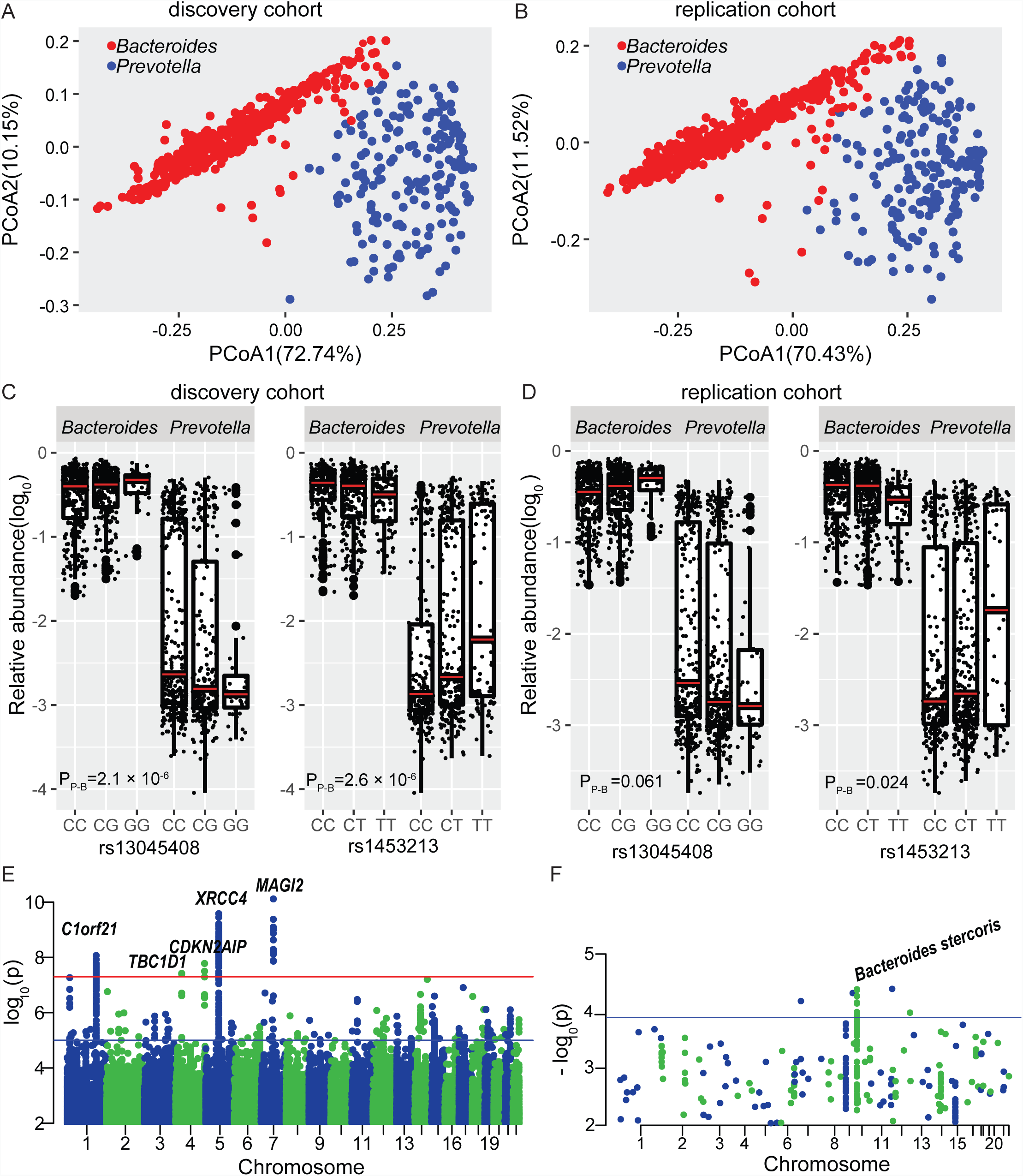
Identifying host genetic variants associated with microbiome enterotypes and principal coordinates (PCoAs, computed using Bray–Curtis dissimilarity). **(A)** The enterotype plot of 618 individuals in discovery cohort. Two clusters were shown with red dots representing *Bacteroides*-dominant enterotype (440 individuals) and blue dots representing *Prevotella*-dominant enterotype (178 individuals). The first 2 principal components (PCoA1 and PCoA2) are shown, with the amount of variation explained are reported for each axe. **(B)** The enterotype plot of 663 individuals in replication cohort. Two clusters were shown with red dots representing Bacteroides-dominant enterotype (473 individuals) and blue dots representing Prevotella-dominant enterotype (190 individuals). **(C)**The minor allele G of SNP rs13045408 at *BTBD3-LINC01722* were positively correlated with *Bacteroides* abundance and negatively correlated with *Prevotella* abundance in discovery cohort. However, SNP rs1453213 at *OXR1* had opposite effects to enterotypes compared with rs13045408 in (B). **(D)** rs13045408 and rs1453213 associated with ‘*Bacteroides*-*Prevotella’* enterotype in replication cohort (*P_P-B_* = 0.061 and 0.024, respectively). **(E)** Manhattan plots of the host genetic variants associated with microbiome β-diversity (computed as Bray–Curtis dissimilarity matrix). The red line represents a genome-wide significant P value (5 × 10^−8^) and blue line represents suggestive P value (10^−5^). Five top loci were marked with gene name. **(F)** The replicated P value in this study for the 380 SNPs previously reported to be significantly associated with the microbiome. Eleven SNPs are successfully replicated at P < 1.3 × 10^−4^ = 0.05/380 (blue line), nine of which were most associated with *Bacteroides stercoris*.

In order to validate these suggestive associations, we sequenced a validation cohort of 663 individuals (metagenomic shotgun sequencing for stool samples to an average of 8.59 ± 2.14 GB (**Fig. S1D**), but 7× whole-genome sequencing for human genome (range from 5× to 12×, **Table S1A**; **Fig. S1C**)). Summary statistics of the covariates was largely similar (**Table S1B**). This replicate cohort comprised of 473 *Bacteroides*-dominant and 190 *Prevotella*-dominant individuals (Fig. 1B). The top 2 associations for the *Bacteroides*-*Prevotella* dichotomy remained (Fig. 1D; **Table S1D**; *P_P-B_* = 0.024 for rs1453213 in *OXR1* and *P_P-B_* = 0.061 for rs13045408 at *BTBD3*-*LINC01722*).

We next investigated associations between genetic variation and microbiome β-diversity. This analysis found five loci with marginal genome-wide significance (*P* < 5 × 10^−8^; Fig. 1E; **Table S1F**). Three SNPs, rs60689247 in *MAGI2*, rs7716962 in *XRCC4* and rs61823500 in *C1orf21*, were located in the intronic region of genes. *MAGI2* related to multiple phenotypes or diseases in the GWAS catalog (MacArthur et al., 2017), including body mass index (BMI), schizophrenia, coronary artery calcification and type 2 diabetes, etc. The protein encoded by *XRCC4* functions together with DNA ligase IV and the DNA-dependent protein kinase in the repair of DNA double-strand breaks. The other two SNPs, rs11732767 and rs1967284 are located in the intergenic regions of *CDKN2AIP* and *TBC1D1*, respectively. *CDKN2AIP* is critical for the DNA damage response and *TBC1D1* is link to Crohn’s disease, lymphocyte count, etc. These are interesting associations, given the increasing incidences of Crohn’s disease and cancer. The association between rs61823500 at *C1orf21* and β-diversity could be replicated both in our replication cohort and German cohort (*P*<0.05)(Wang et al., 2016). However, the three previous studies (Goodrich et al., 2016; Rothschild et al., 2018; Wang et al., 2016) identified a total of 64 SNPs associated with beta-diversity of gut microbiota. Of these, only one SNP was replicated here with nominal significance (*P* = 0.013, **Table S1G**), and none was significant after multiple-test correction. 8 of the 64 SNPs were not found or rare in the Chinese population (MAF < 0.01). The allele frequencies of these 64 SNPs were significantly different between Chinese and European populations (T-test *P*_*difference*_=1.55 × 10^−5^, **Fig. S3**). Besides, 380 SNPs have been previously reported to associate with specific taxa and we were able to replicate 11 of these (*P* < 0.05/380 =1.3 × 10^−4^; Fig. 1F; **Table S1H**), especially in association with *Bacteroides stercoris* (Rothschild et al., 2018). 92 of the 380 loci were not found or rare in Chinese population. In summary, huge population heterogeneity exists, as is known from GWAS studies(Wojcik et al., 2019), and it is necessary to identify Asian-specific host genome-microbiome associations for better understanding genome–microbiome interactions among different ethnicities.

### Heritability estimates underscore genetic contributions to gut microbiome

To assess the influence of host genetics on the gut microbiota, we calculated heritability estimates 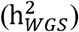 for each specific taxon using whole-genome genetic variants. Among the 510 common taxa with at least 50% occurrence rate of all samples, 39 taxa (7.6%) had a moderate or higher heritability greater than 0.6; 26 taxa are significantly heritable with 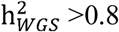 and likelihood-ratio test (LRT) *P* < 0.05 while evaluating with GREML-LDMS (Yang et al., 2015) (Fig. 2A and **Table S2A**). These heritable taxa mainly belong to the Bacteroidetes, Firmicutes and Actinobacteria phyla. *Prevotella* and *Bifidobacterium*, displayed the highest heritability at genus level with 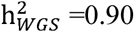 and 0.88, respectively (Fig. 2A). Similarly, their members, species *Prevotella melaninogenica*, *Prevotella veroralis* and *Prevotella bryantii belonging to Prevotella*, species *Bifidobacterium longum* and *Bifidobacterium catenulatum*-*Bifidobacterium pseudocatenulatum* complex belonging to *Bifidobacterium*, all showed high heritability (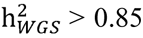, LRT *P* < 0.05). Our results are consistent with the UK twins’ (Xie et al., 2016) and Korean twins’ studies (Lim et al., 2017) which reported that genus *Prevotella* (h = 0.57) and *Bifidobacterium* (h = 0.457) had high heritability, respectively. These were also heritable in the low-depth validation cohort (*Prevotella* 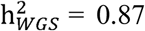; LRT P=0.037, and *Bifidobacterium* 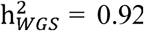; LRT P=0.030, **Table S2A**).

**Figure 2.**
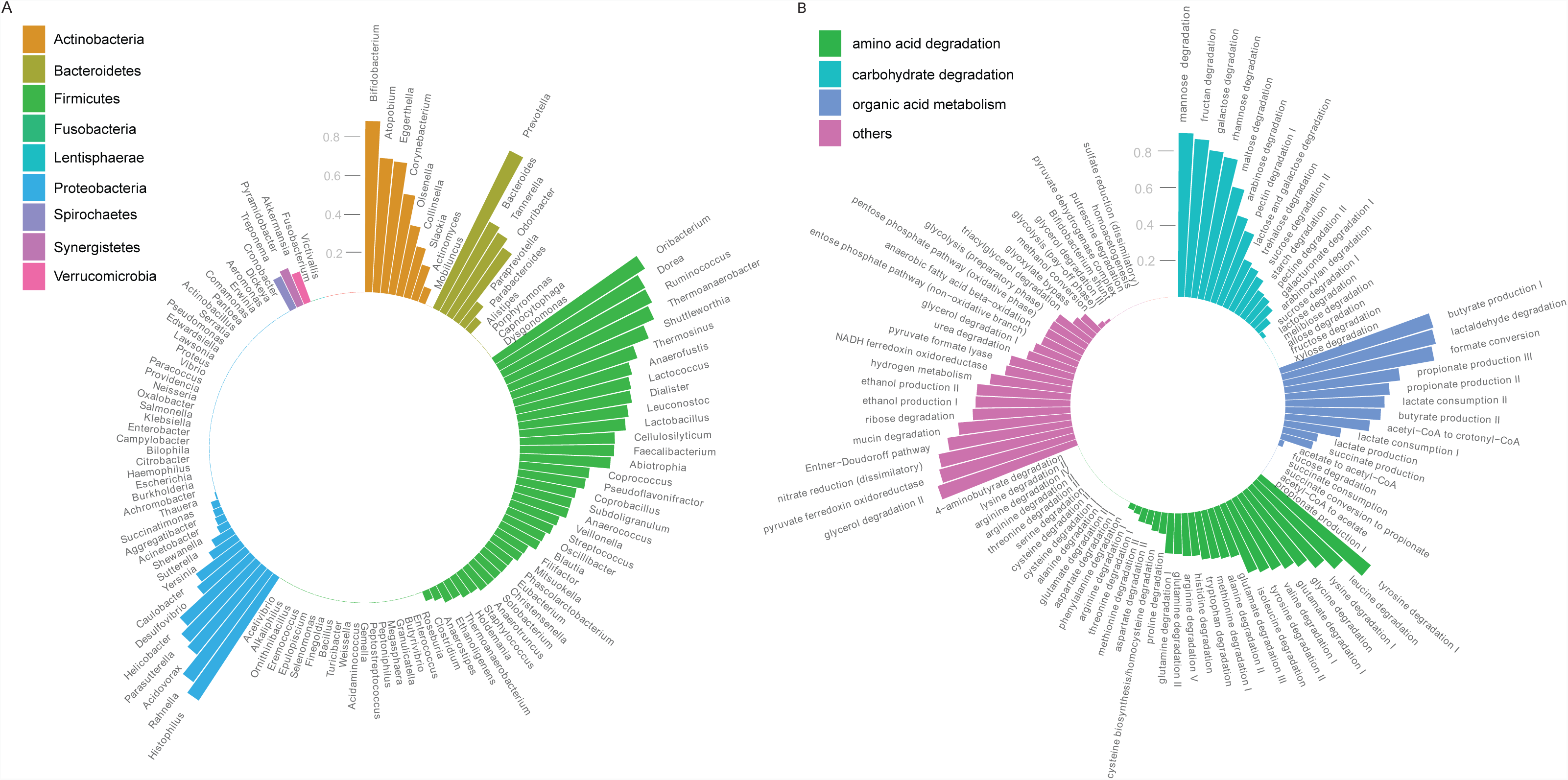
Heritability of gut microbial taxa and its functions. **(A)** The heritability (h2 calculated using GREML-LDMS model in GCTA tool using host whole genome) was plotted as a bar for each genus. Two significantly heritable genura were focus on *Prevotella* and *Bifidobacterium*. More detailed results for heritability of taxa are available in Table S2A. **(B)** Heritability of gut microbial modules (GMMs), which were colored by different metabolic types, including amino acid degradation (red), carbohydrate degradation (green), organic Acid metabolism (blue) and others (purple). Detailed results for heritability of GMMs are available in Table S2C.

Taking advantage of the metagenomic shotgun data, we further evaluated heritability of the functional capacity of the gut microbiome, according to the Kyoto Encyclopedia of Genes and Genomes (KEGG) orthologues (KOs) and gut microbial modules (GMMs) profiles. We identified 4919 common KOs and 98 common GMMs present in 50% or more of the samples. Of which, 152 KOs and 12 GMMs showed high heritability (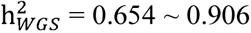, LRT *P* < 0.05, Fig. 2B, **Table S2B,C**). These heritable functional modules included carbohydrate metabolism (mannose degradation, fructan degradation, galactose degradation, rhamnose degradation and maltose degradation), organic acid metabolism (butyrate production, lactaldehyde degradation and formate conversion) and lipid metabolism (glycerol degradation). Among the 152 heritable KOs, 41 KOs were enriched in gut microbiome data of type 2 diabetes (T2D) patients compared to non-diabetic controls (Qin et al., 2012). Of which, aldehyde reductase (K00011) and PTS system, galactitol-specific IIB component (K02774) belonged to the galactose metabolism pathway (ko00052); pyruvate, orthophosphate dikinase (K01006) belonged to the pyruvate metabolism pathway (ko00620). These two KEGG pathways were previously reported to be associated with T2D risk by genome-wide association study (Perry et al., 2009). Furthermore, three KOs, including two T2D-associated KOs K0008 and K01138, and K04075 were confirmed as heritable in the replication cohort (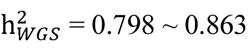, LRT *P* < 0.05; **Table S2B**). We also confirmed one heritable GMM, i.e. butyrate production I (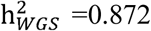, LRT *P* =0.049; **Table S2C**), consistent with twins’ data from UK (Xie et al., 2016). These evidences suggest that some of the heritable KOs in the gut microbiome were determined by host genetic variants and may increase the risk of metabolic diseases such as T2D.

Next, we investigated the genetic variants correlated with functional capacity of the gut microbiome according to GMMs. We found eight loci significantly associated with seven GMMs (*P* < 5 × 10^−8^; **Table S2D**). The strongest association was identified for maltose degradation 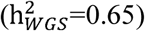 and nine SNPs (*P*=4.5 × 10^−9^) at *SLC41A2* gene which involved in transport of glucose and other sugars, bile salts and organic acids, metal ions and amine compounds. We also found genetic signals for butyrate production 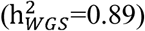 and mucin degradation 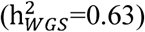. Mucin degradation has been implicated with metabolic regulation, obesity and type 2 diabetes. Three SNPs near *CCR3* gene associated with mucin degradation and *CCR3* gene was reported as enhancer of BMI-related phenotypes and coronary artery disease (*P*<1.0 × 10^−8^) in transcriptome-wide association study (Gusev et al., 2016), and our results suggest that mechanistic investigations of these SNPs should take the gut microbiome into consideration.

### Common variants M-GWAS identifying abundant genetic signals for gut microbiome

To detect associations between gut microbial taxa and specific genetic variants, we at first performed common variants M-GWAS analysis using a linear model for taxa present in over 95% of individuals and a logistic model for zero-inflated taxa (**Table S3A**). We identified 347 significant associations involving 37 loci and 51 bacterial taxa (*P* < 5 × 10^−8^; Fig. 3A, **Table S3B**). The strongest signal (*P* = 1.68 × 10^−9^; **Fig. S4A,B**) was observed for the phylum Actinobacteria and its members, which is consistent with their high heritability (Fig. 2A). Actinobacteria associated with SNP rs62183161 at *LOC150935* gene, which had been reported linked to body composition measurement and energy intake (Comuzzie et al., 2012). Similarly, the other highly heritable taxa, family Prevotellaceae was associated with SNP rs1453123 in *OXR1* (oxidation resistance 1; *P* = 1.58 × 10^−8^; **Fig. S4C,D**) gene, the protein encoded by which controls the sensitivity of neuronal cells to oxidative stress and lacking Oxr1 caused cerebellar neurodegeneration in mice (Oliver et al., 2011). *Eggerthella* abundance was associated with a missense variant rs3749147 in the *GPN1* gene (*P* = 3.2 × 10^−8^). Rs3749147 had been reported to associate with waist circumference - triglycerides in GWAS catalog and *GPN1* gene is linked to oral cavity cancer, palmitoleic acid levels and periodontitis. Seven of the 37 genome-wide significant loci have been reported to associate with traits or diseases in the GWAS catalog (**Table S3C**). We were able to replicate 13 of the 37 associations in the low-depth validation cohort; rs13420238 at *LOC150935* had a *P* = 0.007 with *Bifidobacterium longum*, rs1453123 at *OXR1* had a *P* = 0.006 with Prevotellaceae and rs760646544 near *PLXDC2* have a *P=*0.043 with *Roseburia intestinalis* (Fig. 3B,C,D; **Table S3B**).

**Figure 3.**
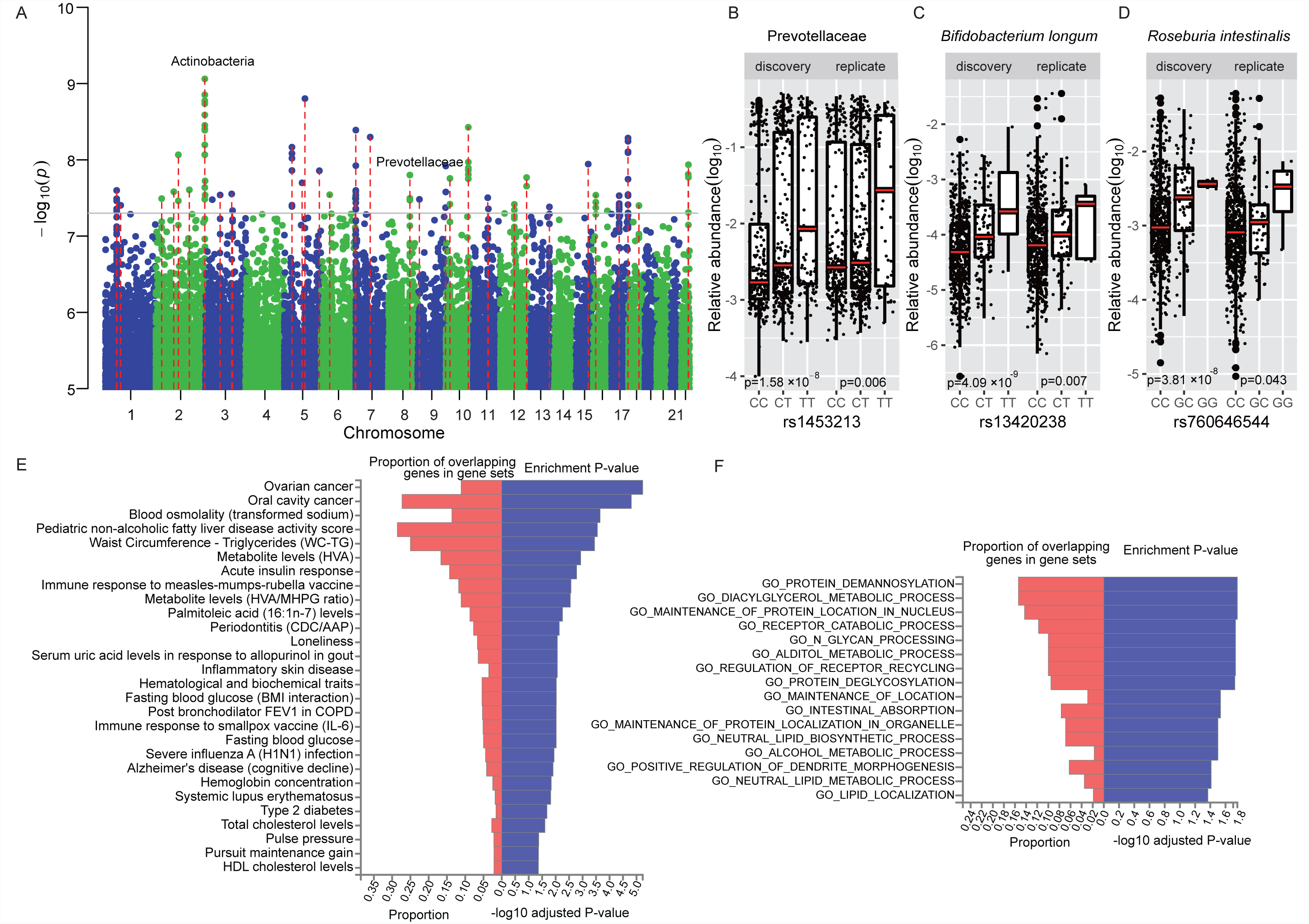
M-GWAS results for the gut microbiome and functional analysis. **(A)** Manhattan plot showing the associations between common variants and taxa, only suggestive associations with *P*<10^−5^ were showed. The gray line represents a genome-wide significant P value 5 × 10^−8^. The red lines showed the 37 loci reaching the genome-wide significance. Two high heritable taxa, Actinobacteria linked to rs13420238 at *LOC150935* and Prevotellaceae linked to rs1453123 at *OXR1*, were marked. **(B)** rs1453123 at *OXR1* associated with Prevotellaceae both in discovery (*P*=1.58×10^−8^) and replication (P = 0.006) cohort. **(C)** rs13420238 at *LOC150935* associated with *Bifidobacterium longum* (Actinobacteria) both in discovery (*P*=4.09×10^−9^) and replication (*P* = 0.007) cohort. **(D)** rs760646544 near *PLXDC2* associated with *Roseburia intestinalis* both in discovery (*P*=3.81×10^−8^) and replication (*P*=0.043) cohort. **(E)** Enriched traits or diseases identified in GWAS catalog using 109 mapped genes from 37 significant loci. **(F)** Enriched KEGG or GO pathways identified for 109 mapped genes from 37 significant loci.

Furthermore, functional annotations using FUMA (Watanabe et al., 2017) tool showed the 37 loci were mapped to 109 genes which were involved in three main traits and diseases in GWAS catalog (MacArthur et al., 2017) (false discovery rate (FDR) adjusted *P* < 0.05, Fig. 3E): (1) metabolism related traits: waist circumference - triglycerides, metabolite levels (homovanillic acid), serum uric acid levels in response to allopurinol in gout, acute insulin response, fasting blood glucose, total cholesterol levels and HDL cholesterol levels; (2) immune-related diseases: Inflammatory skin disease, systemic lupus erythematosus and type 2 diabetes. (3) nervous system related disease: Alzheimer’s disease (cognitive decline) and loneliness. Meanwhile, we perform gene set enrichment analyses and identified 16 significantly enriched KEGG or GO terms after FDR correction (*P* < 0.05, Fig. 3F), including multiple metabolic process such as propanoate, fatty acid, diacylglycerol and alditol metabolism, as well as neutral lipid biosynthetic process.

### Rare variants- and CNVs-based M-GWAS further revealed genetic contributions to gut microbiome

Given the advantages of high-depth WGS and that 85% (36.5M/43M) of the variants were rare (MAF <0.05), we globally considered whether rare variants in any gene and copy number variants contributed to the gut microbiome composition. We tentatively identified 60 associations involving 47 genes and 54 bacterial taxa (P < 2.14 × 10^−6^ = 0.05/27874 for Bonferroni correction of 27,874 individual genes; **Table S3D**), e.g. *PCSK9* gene, a popular target for lowering LDL cholesterol. We evaluated the interaction between proteins encoded by 47 genes by constructing Protein-protein interaction (PPI) networks. We found there were 34 proteins participating in the connected network among the 47 genes (**Fig. S5**). The 34 proteins have more interactions than expected for a random set of proteins of similar size (enrichment *P*= 0.037), indicating functional intersection of the 34 microbiome-associated proteins. KEGG pathway analysis of the 34 genes showed enrichment in 2 main pathways (**Table S3E**), including hsa03030:DNA replication (FDR=0.002) and hsa01100:Metabolic pathways (FDR=0.044).

CNVs-based M-GWAS identified 18 significant CNVs associated with 20 bacterial taxa (P < 6.25 × 10^−6^ = 0.05/8000 for Bonferroni correction of 8K common CNVs with MAF>0.01, **Table S3F**). Of which, 13 CNVs overlaps with reported CNVs recorded in the Database of Genomic Variants (DGV) (MacDonald et al., 2014). Eight CNVs resides within the genes mainly in the intronic region. We identified the species butyrate-producing bacterium SS3/4 associated with a 28.7kb CNV region (chr4:69384168-69412841; *P* = 3.1 × 10^−6^, frequency = 0.03) which located 67 kb downstream of the *UGT2B4* gene. The CNV includes many variants involved in expression quantitative trait loci (eQTL) and regulate the expression of *UGT2B4* in heart tissue and *UGT2B28* in the Esophagus Mucosa and liver (**Table S3G**), consistent with functions of butyrate or other bacterial metabolites in these tissues. Moreover, one SNP rs12505338 and the other three SNPs in the CNV region were associated with serum concentration of stearate (18:0) (*P*=9.3× 10^−5^, **Table S3F**) and glutamate (*P*=7.3× 10^−6^), respectively, by looking for the NHLBI GRASP catalog (Leslie et al., 2014). Thus, associations with the gut microbiome due to rare variants and CNVs also have important functional implications for host physiology.

Common variants-, rare variants- and CNVs-based associations separately explained 8.3%, 11.4% and 4.9% of the microbiome composition, respectively. In addition, the average occurrence frequency of gut microbiome taxa associated with common variants or rare variants were 0.768 and 0.596, respectively (**Tables S3B and S3D**; Wilcoxon test, P=0.008). These results indicate that host rare variants also shaped the gut microbiome, especially for less common members of the community.

### A gene-bacteria axis in gender-differential metabolic and neuronal functions

53.5% of this cohort were female, permitting a comparison between genders. Females showed higher alpha diversity than men (Wilcoxon test, P<0.05; **Fig. S6**), and we identified 32 significantly different taxa between genders in discovery cohort and 27 of which were consistent in replication cohort (FDR q <0.1 using MaAslin; **Table S4A**). Phylum Actinobacteria and its members, including class Actinobacteria, order Bifidobacteriales, family Bifidobacteriaceae, genus *Bifidobacterium* were all significantly enriched in females. However, *Fusobacterium* was significantly enriched in male.

Since the gut microbiome exhibited striking difference between males and females, we performed a gender stratified association analysis between host genetic variants and gut bacteria, the 37 associations were overlapped between genders (**Table S4B**, P<0.05 both in males and in females, and P < 5×10^−8^ in combined results), identical to the combined analysis (Table S3B.). Especially, we identified 33 male-specific (P <5× 10^−8^ in males but P>0.05 in females) and 37 female-specific associations (P <5× 10^−8^ in females but P>0.05 in males) linked to gut bacteria (Fig. 4A and **Table S4C**). We compared the effect sizes of identified variants between genders, and confirmed that all the variants showed a significant difference (*P*_*difference*_ <0.01). Five loci of the 70 associations linked to traits or diseases in GWAS catalog (**Table S4D**). Rs4650205 at *NEGR1-LINC01360* gene significantly associated with the abundance of genus *Acidaminococcus* in males (*P* = 4.87 × 10^−8^) but not in females (*P* = 0.48), and its proxy SNPs (linkage disequilibrium r^2^ > 0.6) were reported linked to multiple nervous-system disorders such as autism spectrum disorder, schizophrenia, depression and migraine by substantial GWAS studies. Female-specific SNP rs61781314 at *LEPR* gene was associated with both genus *Eggerthella* and species *Eggerthella lenta*, and its proxy SNP rs17415296 linked to blood protein levels (*P* = 4 × 10^−229^). *Eggerthella lenta* as an opportunistic pathogen have been reported to underlie human infections and enriched in T2D (Qin et al., 2012), rheumatoid arthritis (RA) (Zhang et al., 2015) and atherosclerotic cardiovascular disease (ACVD) patients (Jie et al., 2017). The protein encoded by *LEPR* is a receptor for leptin (an adipocyte-specific hormone that regulates body weight), and is also involved in the regulation of fat metabolism and pituitary dysfunction. In the low-depth validation cohort (327 males and 336 females), associations including the male-specific association of rs6871146 with *Lactococcus* (β_*male*_=0.42 and *P*_*male*_=0.027; β_*male*_=−0.47 and *P*_*female*_=0.062) and the female-specific association of rs7165633 and *Mobiluncus mulieris* (β_*male*_=0.009 and *P*_*male*_=0.96; β_*female*_=−0.48 and *P*_*female*_=0.008) were replicated. The female-specific association with *Mobiluncus* suggests an intestinal reservoir for the bacterium which is involved in vaginal infections (Onderdonk et al., 2016).

**Figure 4.**
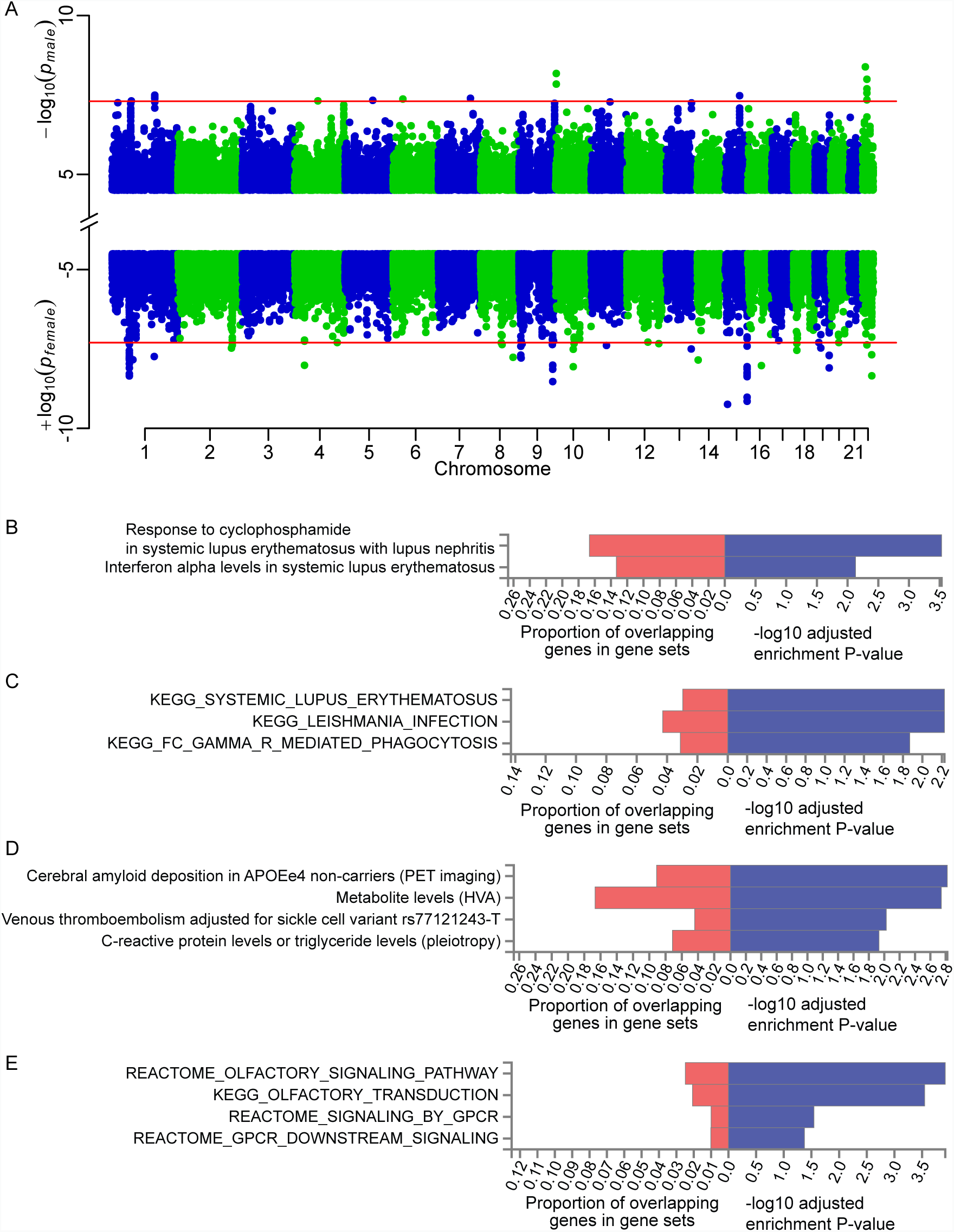
Gender-specific associations and functional analysis. **(A)** Manhattan plot showing the male-specific and female-specific associations for gut microbiome. The red line represents the genome-wide P threshold (5 × 10^−8^). **(B)** Enriched traits or diseases identified using male-specific associations mapped in GWAS catalog. **(C)** Enriched KEGG, REACTOME or GO pathways identified for male-specific associations using FUMA tool. **(D)** Enriched traits or diseases identified using female-specific associations mapped in GWAS catalog. **(E)** Enriched KEGG, REACTOME or GO pathways identified for female-specific associations using FUMA tool.

We investigated the overlapped genes between gender-specific genes and traits or diseases-associated genes in GWAS catalog (Fig. 4B-E), then found that genes located in female-specific loci enriched in four phenotypes, including two metabolic traits, i.e. metabolite levels (HVA) and C-reactive protein levels or triglyceride levels (pleiotropy). Genes located in male-specific loci enriched in mainly the systemic lupus erythematosus related traits. Interestingly, one locus chr19:53772987-53796549 was both male and female-specific locus although associated with different taxa in different genders, and this locus located nearby the gene family *MIR371A-MIR372-MIR373*. WikiPathway analysis showed this gene family related to “miRNAs involved in DNA damage response”, suggesting that gut bacteria may participated in DNA damage response in both genders. Additionally, gender-specific loci were also enriched in the pathway “leptin insulin overlap”, consistent with the association between *LEPR* gene and *Eggerthella lenta* as described above. Moreover, gene set enrichment analysis identified 37 female-specific loci involved in pathways “olfactory signaling” and “signaling by G-protein-coupled receptor (GPCR)”, females had been reported superior to males in olfactory abilities (Brand and Millot, 2001) and had higher levels of G-protein-coupled kinases [GPCR kinase (GRK)] 3 and 5 than male(Bychkov et al., 2011). Male-specific loci related to pathways “systemic lupus erythematosus” and “leishmania Infection”. Taken together, the gut microbiome exhibited differential associations with the human genome in males and females, and might contribute to different metabolic and neuronal functions as well as disease susceptibility.

### M-GWAS helps understand biomarkers from MWAS

We note that our M-GWAS discovered signals for some of the bacteria often reported from metagenome-wide association studies (MWAS) (Wang and Jia, 2016), e.g. three butyrate-producing species, *Roseburia intestinalis*, *Eubacterium rectale* and *Faecalibacterium prausnitzii*, have been associated with healthy controls in MWAS for T2D, ACVD and obesity as well as *Alistipes shahii* associated with lower BMI (Jie et al., 2017; Liu et al., 2017; Qin et al., 2012)(Fig. 5A). We observed the species *Roseburia intestinalis* associated with rs760646544 (a insertion of CTGTT) near *PLXDC2* (*P* = 1.75 × 10^−8^, related to nidogen-1 measurement and diabetic retinopathy in GWAS catalog). Species *Eubacterium rectale* negatively associated with rs1555188 near *PHF21B* in females (*P* = 4.52× 10^−9^) but not in males (*P* = 0.55). Genus *Faecalibacterium* and species *F. prausnitzii* were identified linked to *DYNLL1* gene (*P* = 8.83 × 10^−8^) which included 94 rare variants in gene-based association analysis. *Alistipes shahii* associated with rs72627489 near *SOWAHC* in gender-combined analysis (*P* =8.58 × 10^−9^) and rs914338 near *UNC93A* in male-specific analysis (*P* = 2.40 ×10^−8^). We confirmed the association between *Roseburia intestinalis* and rs760646544 (a insertion of CTGTT) near *PLXDC2* in replicate cohort (*P* = 0.043; Fig. 3D; **Table S3B**). Consistently, the abundance of *R. intestinalis* showed higher correlation in monozygotic compared to dizygotic twins from the United Kingdom(Xie et al., 2016).

**Figure 5.**
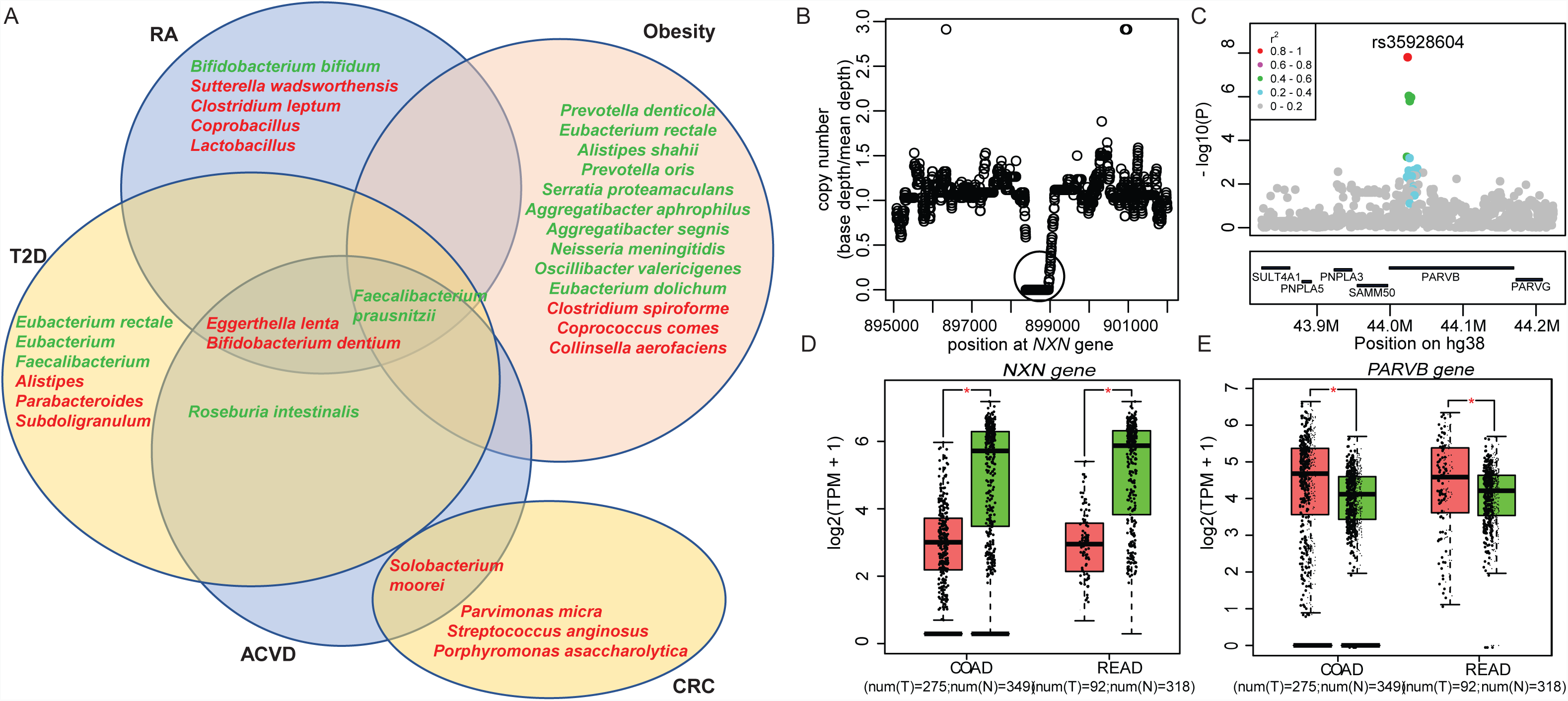
M-GWAS helps understand biomarkers from MWAS. **(A)** Venn diagram showing the bacteria that identified associated with genetic variants and were reported in multiple MWAS studies, including type 2 diabetes (T2D), rheumatoid arthritis (RA), atherosclerotic cardiovascular disease (ACVD), colorectal cancer (CRC) and obesity. Bacteria in overlapping areas indicates it was reported in two or more MWAS studies. The bacteria marked in red represents enrichment in cases, and green represents enrichment in controls. **(B)** A copy loss of 642bp (marked by green circle) in *NXN* gene was detected in Chinese individuals. **(C)** Box plots of the *NXN* gene expression in COAD and READ cases compared with normal tissue samples using GEPIA tool. COAD, colon adenocarcinoma; READ, rectum adenocarcinoma. **(D)** Region plot of the most significant loci for *PARVB* gene. Each point represents a SNP or indel and is colored with the r^2^ value as calculated in this cohort. The lead SNP rs35928604 (P= 1.55 × 10^−8^) is highlighted with red. **(E)** As in **(C)**, box plots of the *PARVB* gene expression in COAD and READ cases compared with normal tissue samples.

*Bifidobacterium dentium*, enriched in RA (Zhang et al., 2015), ACVD as well as schizophrenia patients. CNVs-based M-GWAS analysis identify the association between *Bifidobacterium dentium* and nucleoredoxin (*NXN*) with copy loss of 642 bp (chr17:898377-899018; *P* = 4.11 × 10^−6^; Fig. 5B), and nucleoredoxin 1 as the oxidoreductase protects antioxidant enzymes such as catalase from ROS-induced oxidation in plant cells (Kneeshaw et al., 2017). In addition, *NXN* was significantly high expressed in normal tissue samples compared with colon adenocarcinoma (COAD) and rectum adenocarcinoma (READ) cases (Fig. 5C). Similarly, *Parvimonas micra* is enriched in CRC (Wang and Jia, 2016), its associated host gene *PARVB* (Parvin Beta; rs35928604; *P* = 1.55 × 10^−8^; Fig. 5D) was overexpressed in CRC including COAD and READ (Fig. 5E), which supported the previous study that reported overexpression of *PARVB* correlated significantly with lymph node metastasis and tumor invasion (Bravou et al., 2015). *Bifidobacterium dentium*, *Parvimonas micra* and *Porphyromonas asaccharolytica* are all bacteria found in the human oral cavity that are normally in low abundance in the colon. These findings are consistent with the notion that immune defense are important drivers of host-microbiome co-evolution in addition to metabolism.

## Discussion

The present study is the first complete and comprehensive genome-wide M-GWAS analysis integrating a total of 1295 host whole genome and fecal whole metagenome sequencing to investigate the associations between genetic variants and gut microbiome in Chinese adults. Using common variants-, rare variants-, and CNVs-based association analysis without loosening the p-value cutoff, we identified 37 loci, 47 genes and 20 CNVs significantly associated with gut bacterial taxa, and they additively explained no less than 20% of the microbiome composition. Although study-wise significant associations would be more desirable (P < 1× 10^−10^ for over 500 taxa), consistent with previous M-GWAS from Germany, the Netherlands and Israel(Bonder et al., 2016; Rothschild et al., 2018; Wang et al., 2016), abundant signals were only detected with genome-wide significance. Notably, although insufficient power to detect variants (rare variants and CNVs etc.) in low-depth sequencing data, we still replicated our key findings for ‘enterotypes’, T2D-KOs, common variants’ associations, gender-differential associations, and MWAS markers (Figs. 1, 3, 6, **Table S3B, S4C**). The majority of associations lie in metabolic, neurological and immunological functions, which is particular interesting considering the rapid changes in lifestyle and environmental factors in China and the rising disease incidences. For example, a good portion of our Chinese cohort still harbor *Prevotella* instead of *Bacteroides*, compared with western country (Vangay et al., 2018). To investigate the effect of host genome on enterotype, we identified two suggestive loci explaining 11% of the *Prevotella-Bacteroides* variances. These two tentative associations are not yet genome-wide significant (*P_P-B_* = 2.08 × 10^−6^ and *P*_*P-B*_ = 2.6 × 10^−6^, respectively, using *Prevotella* as cases and *Bacteroides* as controls in logistic regression model), but we feel obliged to report them after validation, given the long-lasting arguments in multiple studies(Costea et al., 2018; Jeffery et al., 2012; Knights et al., 2014) over the concept of ‘enterotypes’. It is intriguing that heterozygous individuals show two clusters of either high or low *Prevotella* (Fig. 1). In addition, we identified heritability and specific loci for *Prevotella* species; the minor allele T of rs1453213 at *OXR1* was consistently correlated with higher abundance of family Prevotellaceae and *Prevotella* species, and higher frequency of allele T in Asian population (f=0.39) than European population (f=0.28) may also explain the enrichment of *Prevotella* in Asian in addition to the diet. More cohorts from developing countries in the future with a higher fraction of *Prevotella*-dominated individuals would help further confirm these results.

Due to the emphasis on diet in early studies and recently on medication, metagenome-wide association studies (MWAS) (Qin et al., 2012; Wang and Jia, 2016) have received even more controversy than GWAS. Besides diet and medication, we also took into account physical activity in this 4D-SZ cohort (Jie et al., 2019a; Jie et al., 2019b). Here we find that fecal biomarkers previously reported by MWAS studies on colorectal cancer and metabolic diseases have some associations with host genetics, while some taxa especially some spore-forming bacteria lacked host genetic associations. With 1 liter of saliva swallowed every day, genetically encoded responses to ectopic presence of oral bacteria in the gut may be a common theme in a number of diseases investigated by MWAS, as has been shown for inflammatory bowel disease (Atarashi et al., 2017).

Gender stratification GWAS could be used to identify novel loci that may have been previously undetected in gender-combined GWAS and had been performed in human complex traits(Khramtsova et al., 2018; Zeng et al., 2018), while none had done it for gut microbiome. Here, we performed the first gender-specific M-GWAS and identified 33 male-specific (involving in inflammation, such as SLE and leishmania infection) and 37 female-specific associations (involving in olfactory signaling and GPCR signaling) linked to gut bacteria by gender-specific analysis, suggesting the importance of discriminating gender in M-GWAS studies and it will help better understand the underlying molecular mechanisms between genders. In summary, our results unveil host genetic influence on the gut microbiome to an unprecedented extent and first reveal the influence of host genome on gender-differential gut bacteria.

## STAR Methods

### KEY RESOURCES TABLE

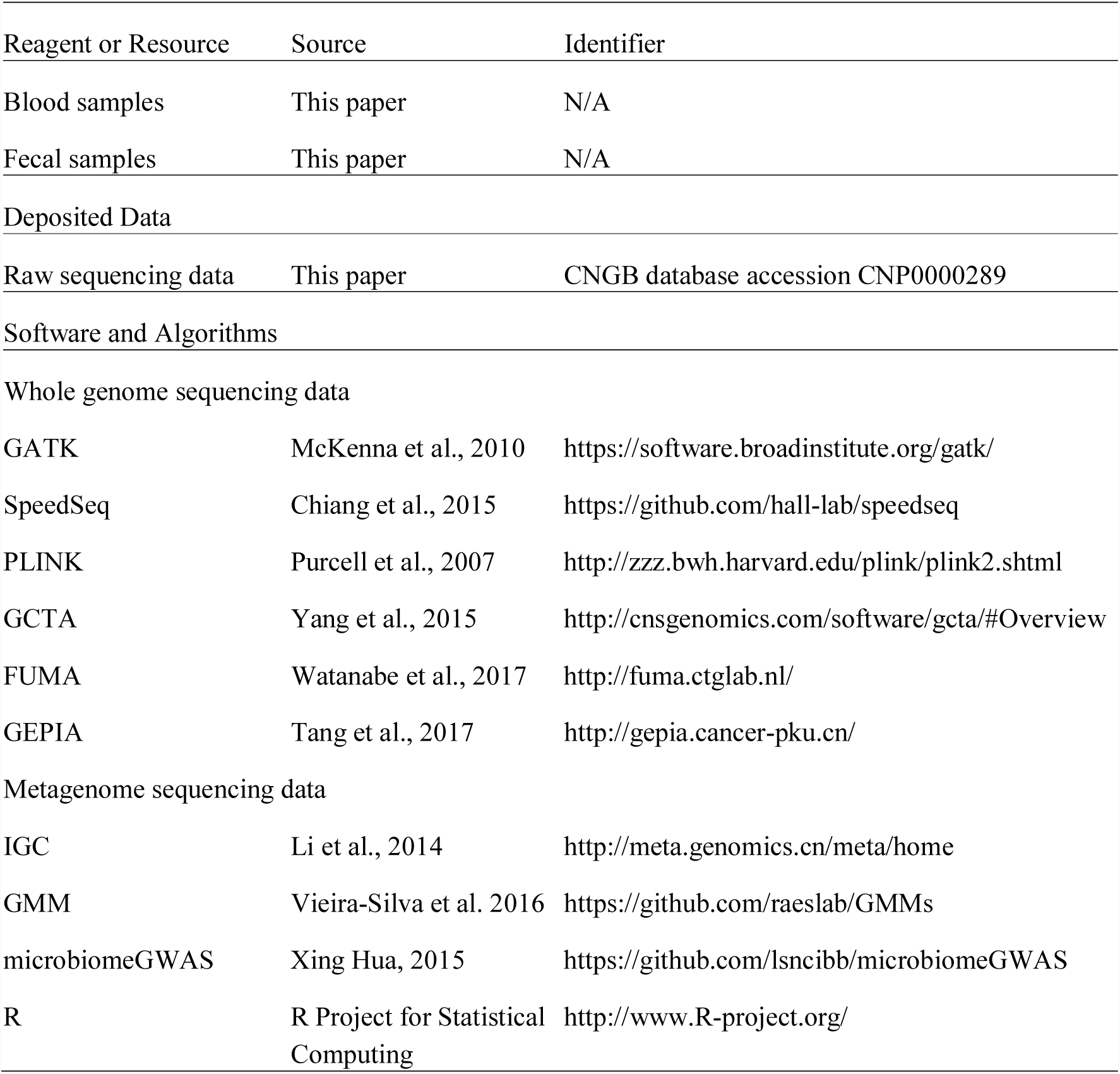

### LEAD CONTACT FOR REAGENT AND RESOURCE SHARING

Further information and requests for resources and reagents should be directed to and will be fulfilled by the Lead Contact, Tao Zhang.

### EXPERIMENTAL MODELS AND SUBJECT DETAILS

#### Cohort descriptions

632 individuals were enlisted in the discovery cohort and 663 individuals were enlisted in the replication cohort, as part of the larger effort of 4D-SZ study(Jie et al., 2019a; Jie et al., 2019b). Questionnaires were collected through a cell phone application. After excluding individuals that were pregnant, taking antibiotics within one month or suffering from diseases, 620 individuals in the discovery cohort and 663 individuals in the replicate cohort were remained. All participants provided blood samples during physical examination. The MGIEasy stool collection kit containing a room temperature stabilizing reagent that preserves metagenomic samples (Han et al., 2018), were also given to the volunteers, who handed in fecal samples on the same morning or the day after. All samples were retrieved from the boxes in front of restrooms and then stored at −80°C before DNA extraction. For blood sample, buffy coat was isolated and DNA was extracted using HiPure Blood DNA Mini Kit (Magen, Cat. no. D3111) according to the manufacturer’s protocol. Feces were collected by MGIEasy and stool DNA was extracted in accordance with the MetaHIT protocol(Qin et al., 2012) as described previously. The DNA concentrations from blood and stool samples were estimated by Qubit (Invitrogen). 200 ng of input DNA from blood and stool samples were used for library formation and then processed for single-end 100bp sequencing on BGISEQ-500 platform(Fang et al., 2018).

The study was approved by the Institutional Review Boards (IRB) at BGI-Shenzhen, and all participants provided written informed consent at enrolment.

### METHOD DETAILS

#### High-depth WGS alignment and SNP/indel calling in discovery cohort

Whole-genome reads were aligned to latest reference human genome GRCh38/hg38 with BWA(Li and Durbin, 2009) (version 0.7.15) with default parameters. The reads consisting of base quality <5 or containing adaptor sequencing were filtered out. The alignments were indexed in the BAM format using Samtools (Li et al., 2009a) (version 0.1.18) and PCR duplicates were marked for downstream filtering using Picardtools (version 1.62). The Genome Analysis Toolkit’s (GATK(McKenna et al., 2010), version 3.8) BaseRecalibrator created recalibration tables to screen known SNPs and INDELs in the BAM files from dbSNP (version 150). GATKlite (v2.2.15) was used for subsequent base quality recalibration and removal of read pairs with improperly aligned segments as determined by Stampy. GATK’s HaplotypeCaller were used for variant discovery. GVCFs containing SNVs and Indels from GATK HaplotypeCaller were combined (CombineGVCFs), genotyped (GenotypeGVCFs), variant score recalibrated (VariantRecalibrator) and filtered (ApplyRecalibration). During the GATK VariantRecalibrator process, we took our variants as inputs and used four standard SNP sets to train the model: (1) HapMap3.3 SNPs; (2) dbSNP build 150 SNPs; (3) 1000 Genomes Project SNPs from Omni 2.5 chip; and (4) 1000G phase1 high confidence SNPs. The sensitivity threshold of 99.9% to SNPs and 99% to indels were applied for variant selection after optimizing for Transition to Transversion (TiTv) ratios using the GATK ApplyRecalibration command. After applying the recalibration, there are 43,342,216 raw variants left, including 38 million SNPs, 5 million indels.

We applied a conservative inclusion threshold for variants: (i) mean depth >8×; (ii) Hardy-Weinberg equilibrium (HWE) *P* > 10^−4^; and (iii) genotype calling rate > 98%. We demanded samples to meet these criteria: (i) mean sequencing depth > 20×; (ii) variant call rate > 98%; (iii) no population stratification by performing principal components analysis (PCA) analysis implemented in PLINK(Purcell et al., 2007) (version 1.07) and (iv) excluding related individuals by calculating pairwise identity by descent (IBD, Pi-hat threshold 0.1875) in PLINK. Only 2 samples were removed in quality control filtering and 618 individuals entered into subsequent analysis.

#### CNV calling

The CNV call set were produced using the SpeedSeq(Chiang et al., 2015) pipeline, followed by the svtools package (v0.2.0; https://github.com/hall-lab/svtools). Briefly, speedseq sv, which comprises LUMPY for SV calling based on discordant pairs and split-reads; svtyper for SV genotyping; and cnvnator for read-depth based CNV detection; was run on each sample individually. The individual-level calls were sorted and merged using svtools lmerge, and then each sample was re-genotyped and copy number annotated at all variant positions using svtools genotype and copynumber, and pasted into a single cohort-level VCF. For filtering, inversion calls and adjacencies (i.e. BNDs) were excluded. The CNV was defined as known in the Database of Genomic Variants (DGV) (MacDonald et al., 2014) (http://projects.tcag.ca/variation) if it had 70% region overlapped with one CNV in DGV.

#### Low-depth WGS alignment and SNP/indel calling in replicate cohort

We used BWA to align the whole genome reads to GRCh38/hg38 and used GATK to perform variants calling by applying the same pipelines for high-depth WGS data. After finishing the GenotypeGVCFs process, we got 29906793 raw variants. A more stringent process in the GATK VariantRecalibrator stage compared with high-depth WGS was then used, as are recommended for low-coverage whole-genome data, to filter the uncertain genotype calls and keep only high-quality variants. Specifically, we excluded SNPs with low mapping quality (Q<20) and SNPs with low depth (DP<3). Further, we kept variants with less than 30% missing information, leaving 779521 highly reliable variants. All these high-quality variants were then imputed using BEAGLE 6(Browning et al., 2018) with 618 high-depth WGS dataset as reference panel. We retained only variants with imputation info. > 0.7 and got 5318809 imputed variants. Finally, we further filtered this set to keep variants with Hardy-Weinberg equilibrium P > 10^−5^ and genotype calling rate > 90%, yielding 5249443 variants for subsequent analysis.

To evaluate the data quality, we sequenced 27 samples with both high-depth and low-depth WGS data and then compared the 5318809 variants between them for each individual. The average genotype concordance was 98.66% (**Table S1I**).

#### Metagenomic sequencing and profiling

The high-quality metagenomic sequencing reads were aligned to hg38 using SOAP2(Li et al., 2009b) (version 2.22; identity ≥ 0.9) to remove human reads. The gene profiles were generated by aligning high-quality sequencing reads to the integrated gene catalog (IGC)(Li et al., 2014) by using SOAP2 (identity ≥ 0.95) as previously described. The relative abundance profiles of phylum, order, family, class, genera, species and Kyoto Encyclopaedia of Genes and Genomes (KEGG)(Kanehisa et al., 2014) orthologous groups (KOs) were determined from the gene abundances. To eliminate the influence of sequencing amount in comparison analyses, we downsized the unique IGC mapped reads to 20 million for each sample. The relative abundance profiles of gene, phylum, order, family, class, genus, species and KOs were determined accordingly using the downsized mapped reads per sample.

GMMs (gut metabolic modules) reflect bacterial and archaeal metabolism specific to the human gut, with a focus on anaerobic fermentation processes(Vieira-Silva et al., 2016). The current set of 103 GMMs was built through an extensive review of the literature and metabolic databases, inclusive of MetaCyc(Caspi et al., 2014) and KEGG, followed by expert curation and delineation of modules and alternative pathways. And we identified 98 common GMMs present in 50% or more of the samples.

#### Covariates used in this study

As part of the 4D-SZ cohort, all participants in this study had records of multi-omics data, including anthropometric measurement, stool form, defecation frequency, diet, lifestyle, blood parameters, hormone, etc. (Jie et al., 2019b). We tested for associations between these environmental factors and microbiome β-diversity at the genus level. The effect size and significance of the mentioned variables were estimated using ‘envfit’ function in vegan (R 3.2.5, vegan package 2.4-4). Gender, BMI and defecation frequency were identified to be the strongest factors to explain gut microbiome composition. They accounted for 3.79%, 2.14% and 1.79% of the microbiome variance, respectively. In addition, given the effects of diet and lifestyles on specific taxa, we finally included age, gender, BMI, defecation frequency, stool form, 12 diet and lifestyle factors, as well as the top four principal components (PCs) as covariates for all subsequent M-GWAS analysis (**Table S1B**).

#### Enterotype analysis

The enterotypes analysis was performed using genus-level gene abundance data according to the Dirichlet multinomial mixtures (DMM) based clustering approach(Ding and Schloss, 2014; Holmes et al., 2012) and two enterotypes were identified among the 618 healthy Chinese individuals in discovery cohort, including *Bacteroides* (enterotype 1, n=440) and *Prevotella* (enterotype 2, n=178). Using the same method, this replicate cohort comprised of 473 *Bacteroides*-dominant and 190 *Prevotella*-dominant individuals. We used logistic model implemented in PLINK to run a GWAS for genetic variation and the enterotype phenotype (ie. *Bacteroides* and *Prevotella*; dichotomous trait). We estimated the proportion of enterotypes’ variance explained by top two loci using the restricted maximum likelihood (REML) method implemented in GCTA.

#### Association analysis for microbiome β-diversity

The microbiome β-diversity (between-sample diversity) based on genus-level abundance data were generated using the ‘vegdist’ function (Bray–Curtis dissimilarities). Then, we performed principal coordinates analysis (PCoA) based on the calculated beta-diversity dissimilarities using the ‘capscale’ function in ‘vegan’. The associations between genetic variants and microbiome β-diversity was performed using microbiomeGWAS(Xing Hua, 2015) tool.

#### WGS-based heritability for gut microbiome

Heritability was estimated with the GREML-LDMS(Yang et al., 2015) method implemented in GCTA(Yang et al., 2011) software. The GREML-LDMS method is proposed to estimate heritability using whole genome sequence (WGS) data. The method is unbiased and corrects for the LD bias in the estimated SNP-based heritability. The analysis involves four steps: 1) calculating segment-based LD score; 2) stratifying SNPs based on the segment-based LD score (this is done in R); 3) computing genotypic relatedness matrix (GRM) using the stratified SNPs; 4) performing REML analysis using the multiple GRMs. Therefore, the heritability 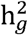 is then the fraction of the variance accounted for by the genetics and can be formulated as: 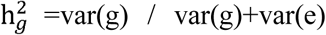, where var(g) and var(e) are the genetic and residual variance components estimated by the REML approach.

In the present study, we estimate heritability 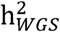 for the relative abundances of each taxon, KO and GMM conditioned on age, gender, BMI, defecation frequency, stool form, self-reported diet, lifestyle factors and top four PCs. We then defined the taxa, KOs and GMMs with LRT P<0.05 in GREML-LDMS as significantly heritable.

#### Genome-wide Association analysis for gut bacteria

We tested the associations between host genetics and gut bacteria using linear or logistic model based on the abundance of gut bacteria. The abundance of bacteria appeared in over 95% of individuals was transformed by the natural logarithm and the outlier individual who was located away from its mean by more than five standard deviations was removed, so the abundance of bacteria could be treated as quantitative trait. Otherwise, we dichotomized bacteria into presence/absence patterns to prevent zero inflation, then the abundance of bacteria could be treated as dichotomous trait. Next, for the common variants with MAF > 5%, we performed a standard single variant (SNP/indel)-based GWAS analysis via PLINK using a linear model for quantitative trait or a logistic model for dichotomous trait, a threshold of *P* < 5 × 10^−8^ was used for genome-wide significance. We used the same methods for CNVs-based association analysis and set a significance threshold at *P* < 6.25 × 10^−6^ accounting for 8,006 common CNVs (MAF > 1%). For rare variants-based association analysis, we applied the Sequence Kernel Association Test(Wu et al., 2011) (SKAT) to the rare variants (MAF ≤ 5%) for each gene. Gene regions were annotated using the RefSeq(Pruitt et al., 2012) database with a total of 27,874 genes. We only included the genes which had five or more rare variants (as recommended by the SKAT authors) for testing; 22,015 genes satisfied this requirement. Associations were considered significant with p < 2.14 × 10^−6^ (equal to 0.05 /22,015). When testing all the association analysis, we adjusted for gender, BMI, defecation frequency, stool form, self-reported diet, lifestyle factors and the first four PCs.

The additive effect of the significant loci from this analysis was then determined using redundancy analysis based on genus-level composition (‘rda’ in the ‘vegan’ package) and the ‘ordiR2step’ function in the ‘vegan’ package in R. To quantify the fraction of microbiome variance that could be inferred from gene-based analysis (actually rare variants), we first selected 200 top-ranking rare variants (not in linkage equilibrium) according to their association with taxa, then performed a greedy stepwise algorithm, in which at each iteration we added the most significant variant to the inferred variant sets added in previous iterations. Before adding each variant, we performed 1000 permutation tests and verified that its contribution was greater than in at least 50% of these permutations. If not, we stopped the algorithm. In each permutation we assigned the top 200 rare variants of each individual to a random individual, and then reran the entire analysis. Finally, 37 loci from common variants-based association analysis, 76 loci from rare variants-based association analysis and 22 loci from CNVs-based association analysis were used to infer the variance, respectively, and then in combination.

#### Functional annotation of significant loci

Genome-wide significant loci identified in M-GWAS analysis were mapped to genes using SNP2GENE in FUMA(Watanabe et al., 2017) (http://fuma.ctglab.nl/). We first converted the loci positions from hg38 to hg19, then used the positional mapping method and maps variants to genes based on physical distance within a 20kb window. Mapped genes were further investigated using the GENE2FUNC procedure, which provides hypergeometric tests of enrichment of the list of mapped genes in 53 GTEx tissue-specific gene expression sets, 7,246 MSigDB gene sets, and 2,195 GWAS catalog(MacArthur et al., 2017) gene sets. Specifically, the background genes in the GENE2FUNC is there for the N which is supposed to be all the genes we considered to select a set of interested genes n. And we have a tested gene set with m genes. The number of overlapped genes between n and m is x. Therefore, the null hypothesis is finding x genes given N, n and m is not more than expected. For example, the GWAS catalog gene sets were defined by extracting genes for each trait from the GWAS catalog. Using the GENE2FUNC procedure, we examined whether the mapped genes enriched in some specific diseases or traits in GWAS catalog as well as whether enriched in specific GO, KEGG et al. The significant results were selected if Bonferroni-corrected *P*<0.05.

#### PPI network analysis

The PPI network was constructed with the Search Tool for Retrieval of Interacting Genes/Proteins (STRING(Szklarczyk et al., 2019), https://string-db.org/cgi/input.pl/). Given a list of the proteins as input, STRING can search for their neighbor interactors, the proteins that have direct interactions with the inputted proteins; then STRING can generate the PPI network consisting of all these proteins and all the interactions between them. We first constructed the PPI network with the 47 significant genes as input, the network displayed on the webpage was gathered into two main clusters and then exported as a high-resolution bitmap. Meanwhile, we got the KEGG pathway enrichment results which were used to characterize the biological importance of the clusters.

#### Gender-specific GWAS analysis for microbiome

We compared the difference of diversity and microbiota composition between genders. Diversity was calculated for Shannon index based on genus-level relative abundance of microbial taxa. Pairwise comparisons were performed using non-parametric test (Wilcoxon test). The multivariate association with linear models (MaAsLin)(Morgan et al., 2012) package was used to identify the differentially abundant taxa between genders. Only taxa with q values <0.05 are identified as significantly enriched in males or females.

We performed gender-specific GWAS analysis in male and female separately using the same methods as described in the microbiome genome-wide association analysis. Male-specific variants were identified as (i) significantly associated with taxa in male (*P*_*male*_ < 5 × 10^−8^) and not significant in female (*P*_*female*_ > 0.05), and (ii) had nominal significant gender difference (testing P value for difference in gender-specific effect size estimated by beta value, *P*_*difference*_ < 0.01). Female-specific variants were identified as (i) significantly associated with taxa in female (*P*_*female*_ < 5 × 10^−8^) and not significant in male (*P*_*male*_ > 0.05), and (ii) had nominal significant gender-difference (*P*_*difference*_ < 0.01, as explained below).

For each variant (SNP/indel/CNV) and for the phenotype (relative abundance of taxa), we computed P values (*P*_*difference*_) testing for difference between the male-specific and female-specific beta-estimates b_male_ and b_female_ using the T-statistic

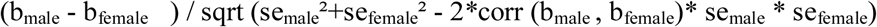

with se_male_ and se_female_ being the standard errors of b_male_ or b_female_. The correlation between the gender-specific beta estimates was computed as the Spearman rank correlation coefficient across all variants for each phenotype.

#### Gene expression and differential analysis

We used GEPIA(Tang et al., 2017) (Gene Expression Profiling Interactive Analysis), a web-based tool to deliver fast and customizable functionalities based on the Cancer Genome Atlas (TCGA) and Genotype-Tissue Expression (GTEx) data. We performed the differential expression analysis for genes *NXN* and *PARVB* across colon adenocarcinoma (COAD) and rectum adenocarcinoma (READ) types compared with paired normal samples, respectively. We choose log2(TPM + 1) transformed expression data for plotting. We used ANOVA method for differential analysis. Genes with higher |log_2_FC| >0.2 and p values <0.05 are considered differentially expressed genes.

## DATA AND SOFTWARE AVAILABILITY

The data in this study have been deposited to the CNGB Nucleotide Sequence Archive (CNSA: https://db.cngb.org/cnsa; accession number CNP0000289).

## Acknowledgments

We are very grateful to colleagues at BGI-Shenzhen for sample collection, DNA extraction, library construction, sequencing, and discussions.

## Author contributions

H.J. and T.Z. conceived and organized this study. J.W. initiated the overall health project. X.X., H.Y. and Y.H. performed the sample collection and questionnaire collection. X.M.L., T.Z., X.T., R.G., Y.Z., X.L., Y.Z. and Y.H. generated and processed the whole genome data. S.T., H.Z., J.Z., Q.D., D.W., L.X. and K.K. generated and processed the metagenome data. X.M.L., S.T., and S.B. performed the bioinformatic analyses, X.M.L. and H.J. wrote the manuscript. All authors contributed to data and texts in this manuscript.

