## Supplemental Figures for "M-GWAS for the gut microbiome in Chinese adults illuminates on complex diseases"

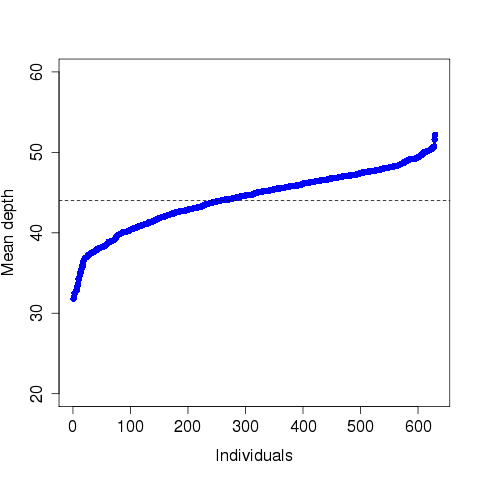

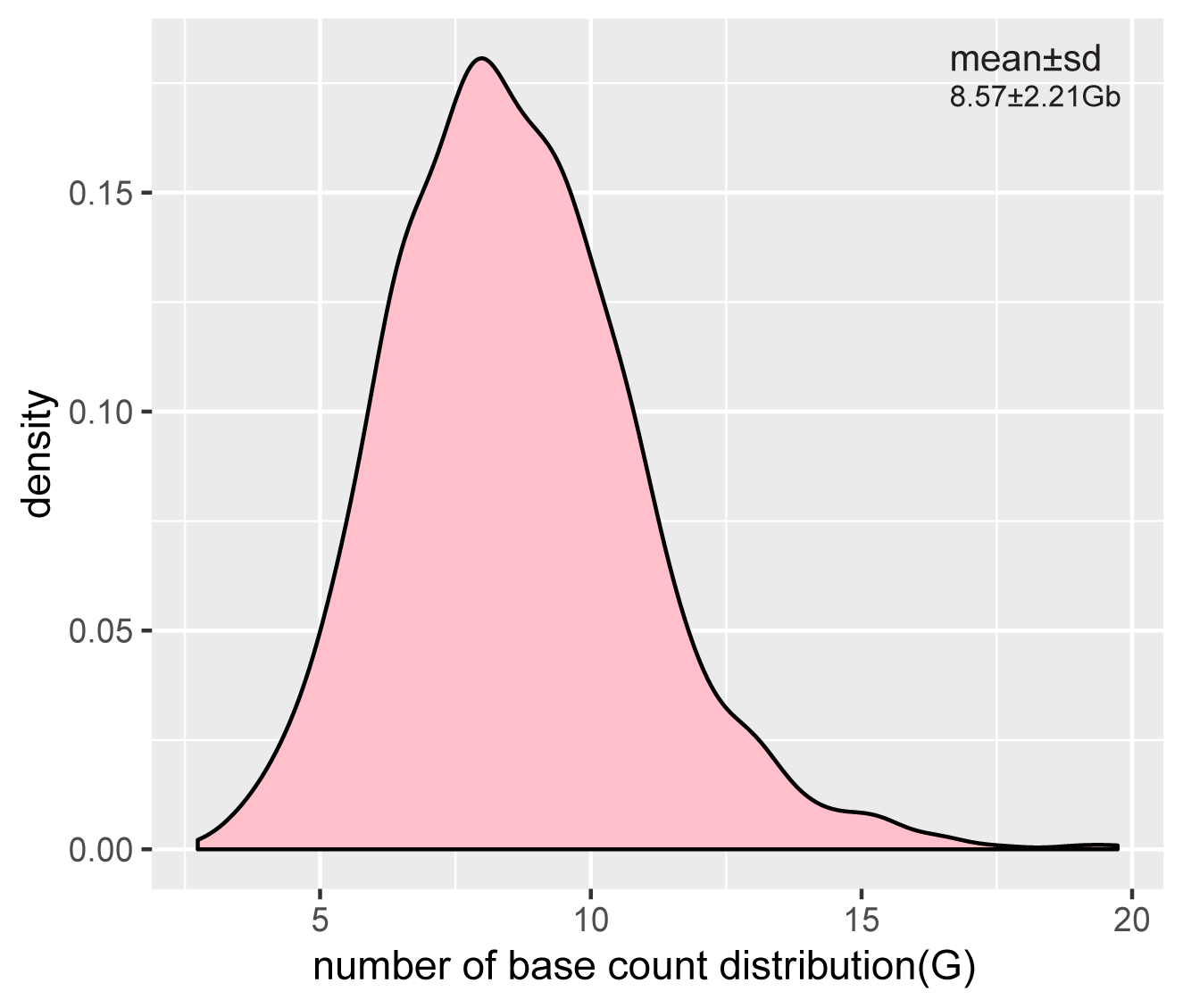

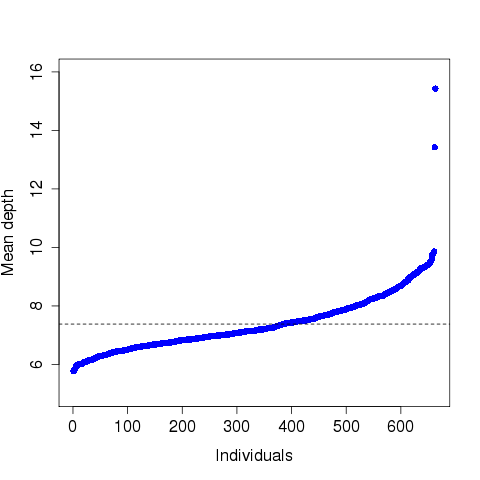

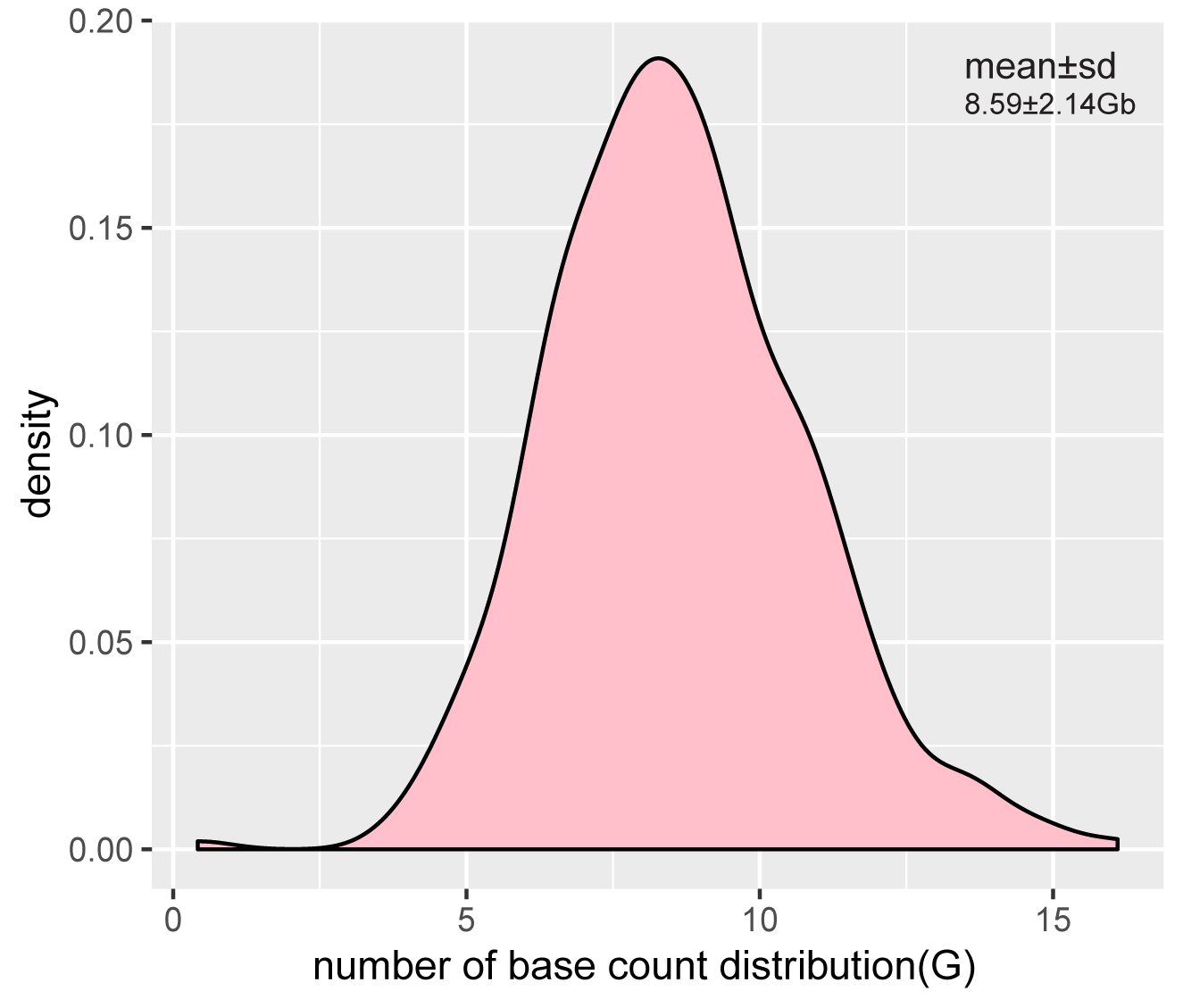

**D**

**C**

**A**

**B**

**Figure S1. Whole-genome sequencing (WGS) and metagenome sequencing data production. (A)** Depth distribution of 632 high-depth WGS samples in discovery cohort. The mean depth is 44×. **(B)** Metagenome sequencing at an average of 8.57±2.21 Gb per sample in discovery cohort. **(C)** Depth distribution of 663 low-depth WGS samples in replication cohort. The mean depth is 7×. **(D)** Metagenome sequencing at an average of 8.59±2.14 Gb per sample in replication cohort.

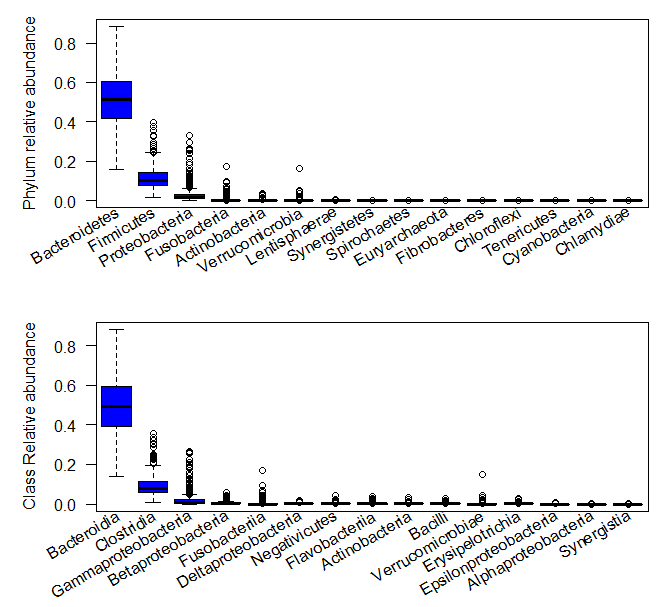

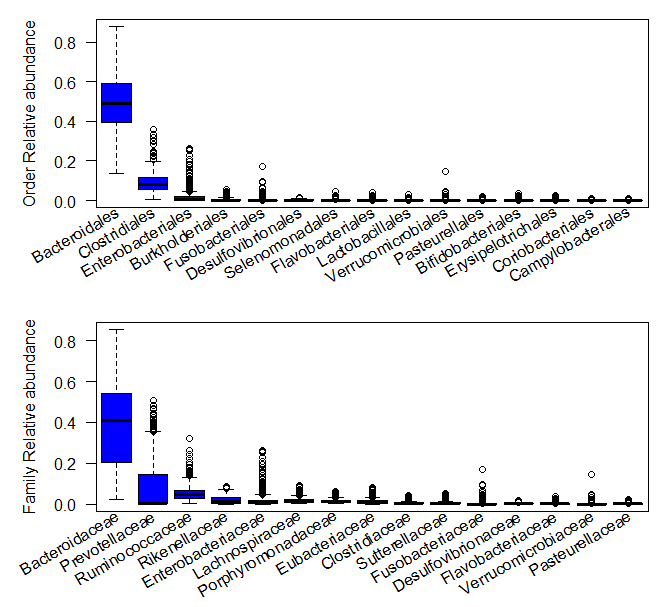

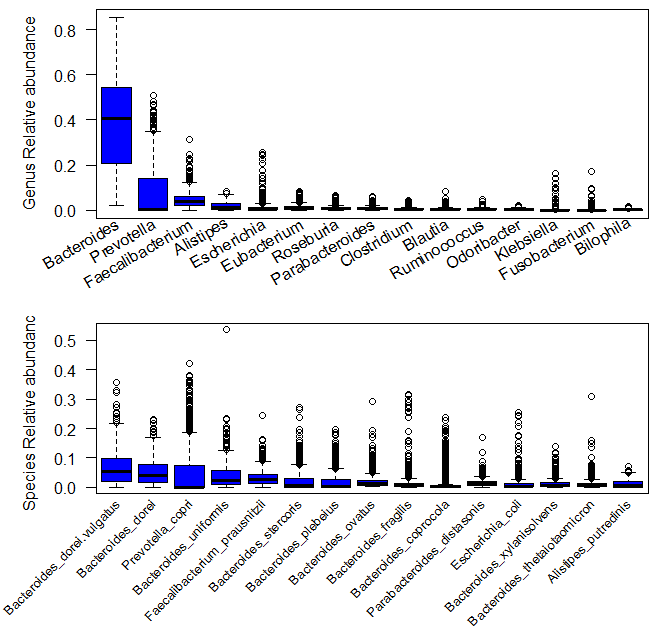

**Figure S2. Box and whisker plot of the top 15 taxa with high relative abundance from phylum to species level in the Chinese cohort.** Boxes represent the interquartile range (IQR) between first and third quartiles and the line inside represents the median. Whiskers denote the lowest and highest values within 1.5 × IQR from the first and third quartiles, respectively. Circles represent outliers beyond the whiskers**.**

|  |  |
| --- | --- |
| 0.345949 | 0.653082 |
| 0.346482 | 0.653082 |
| 0.202559 | 0.43 |
| 0.0463753 | 0.29 |
| 0.479744 | 0.57 |
| 0.208955 | 0.552684 |
| 0.397436 | 0.457256 |
| 0.19 | 0.23 |
| 0.183902 | 0.153082 |
| 0.455 | 0.47 |
| 0.126333 | 0.178926 |
| 0.286247 | 0.35 |
| 0.0175906 | 0.15 |
| 0.493597 | 0.176938 |
| 0.247868 | 0.143141 |
| 0.388593 | 0.224652 |
| 0.273987 | 0.482107 |
| 0.0916844 | 0.54175 |
| 0.152452 | 0.191849 |
| 0.152985 | 0.19 |
| 0.152452 | 0.18 |
| 0.397 | 0.54 |
| 0.3758 | 0.79 |
| 0.286247 | 0.12 |
| 0.433369 | 0.176938 |
| 0.146588 | 0.190855 |
| 0.147655 | 0.191849 |
| 0.147122 | 0.191849 |
| 0.147655 | 0.190855 |
| 0.147655 | 0.191849 |
| 0.147122 | 0.190855 |
| 0.147655 | 0.190855 |
| 0.147655 | 0.190855 |
| 0.147655 | 0.190855 |
| 0.147655 | 0.191849 |
| 0.188166 | 0.42 |
| 0.146588 | 0.175944 |
| 0.146588 | 0.189861 |
| 0.146588 | 0.176938 |
| 0.183369 | 0.196819 |
| 0.133262 | 0.15507 |
| 0.182303 | 0.261431 |
| 0.118337 | 0.154076 |
| 0.0804904 | 0.13 |
| 0.0826226 | 0.128231 |
| 0.032516 | 0.16 |
| 0.0831557 | 0.141153 |
| 0.00319829 | 0.11 |
| 0.00479744 | 0.1 |
| 0.00458635 | 0.16 |
| 0.0095661 | 0.11 |
| 0 | 0.52 |
| 0 | 0.52 |
| 0.000533049 | 0.47 |
| 0 | 0.66 |

**Figure S3. The alleles frequencies difference of reported 64 loci associated with beta-diversity between two populations.** The European population represents the EUR population in 1000genome phase3, and the alleles frequencies were acquired by searching in Haploreg website (https://pubs.broadinstitute.org/mammals/haploreg/haploreg.php).

**B**

**A**

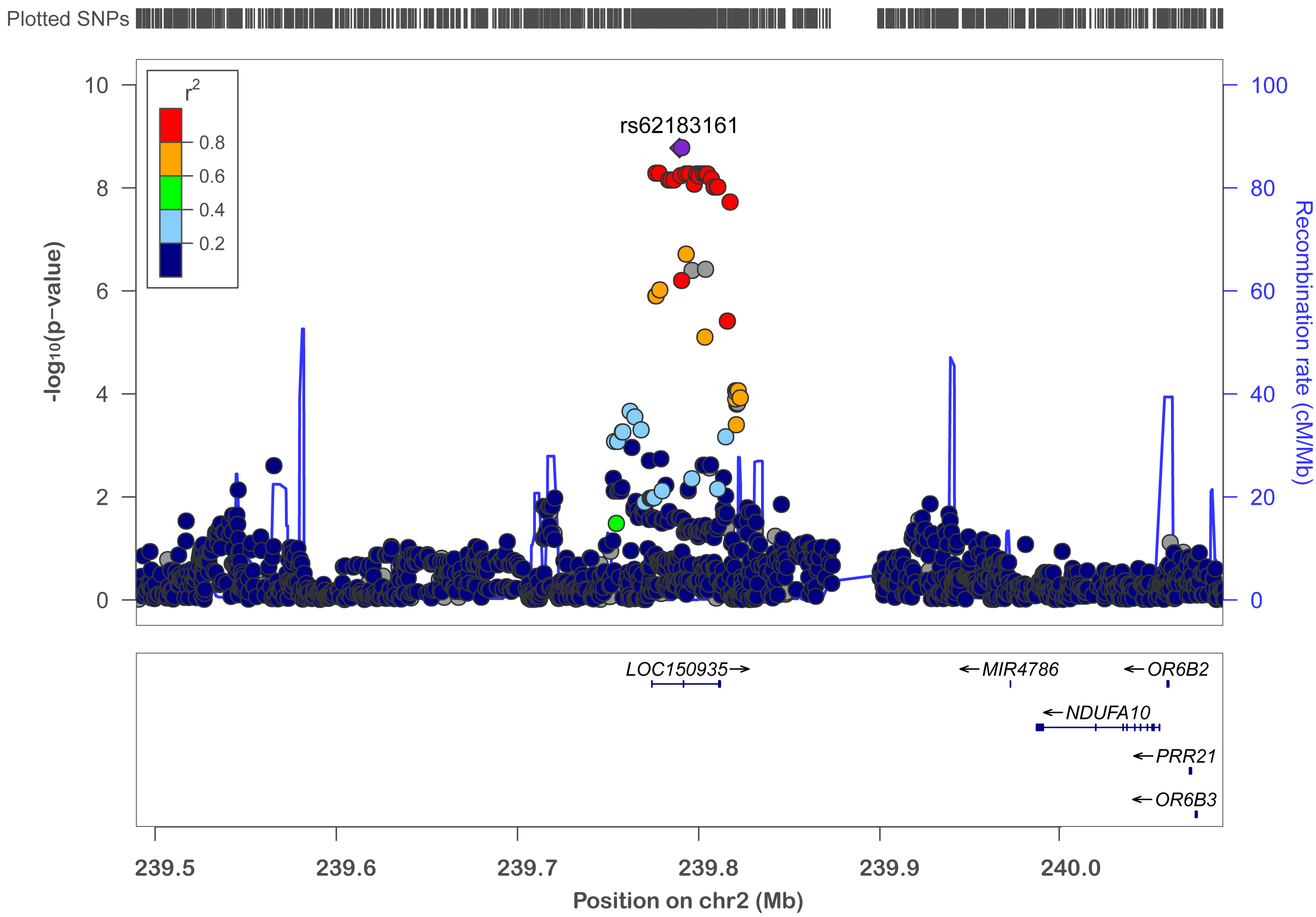

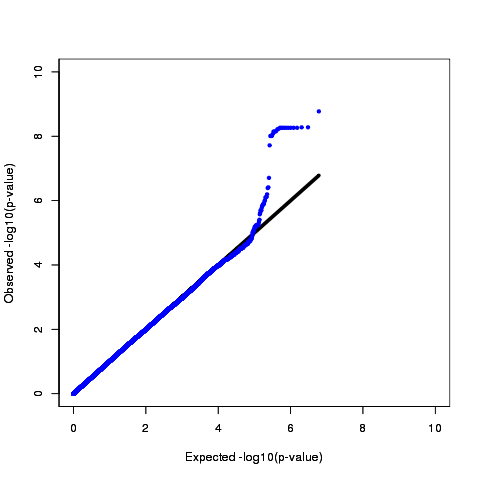

**D**

**C**

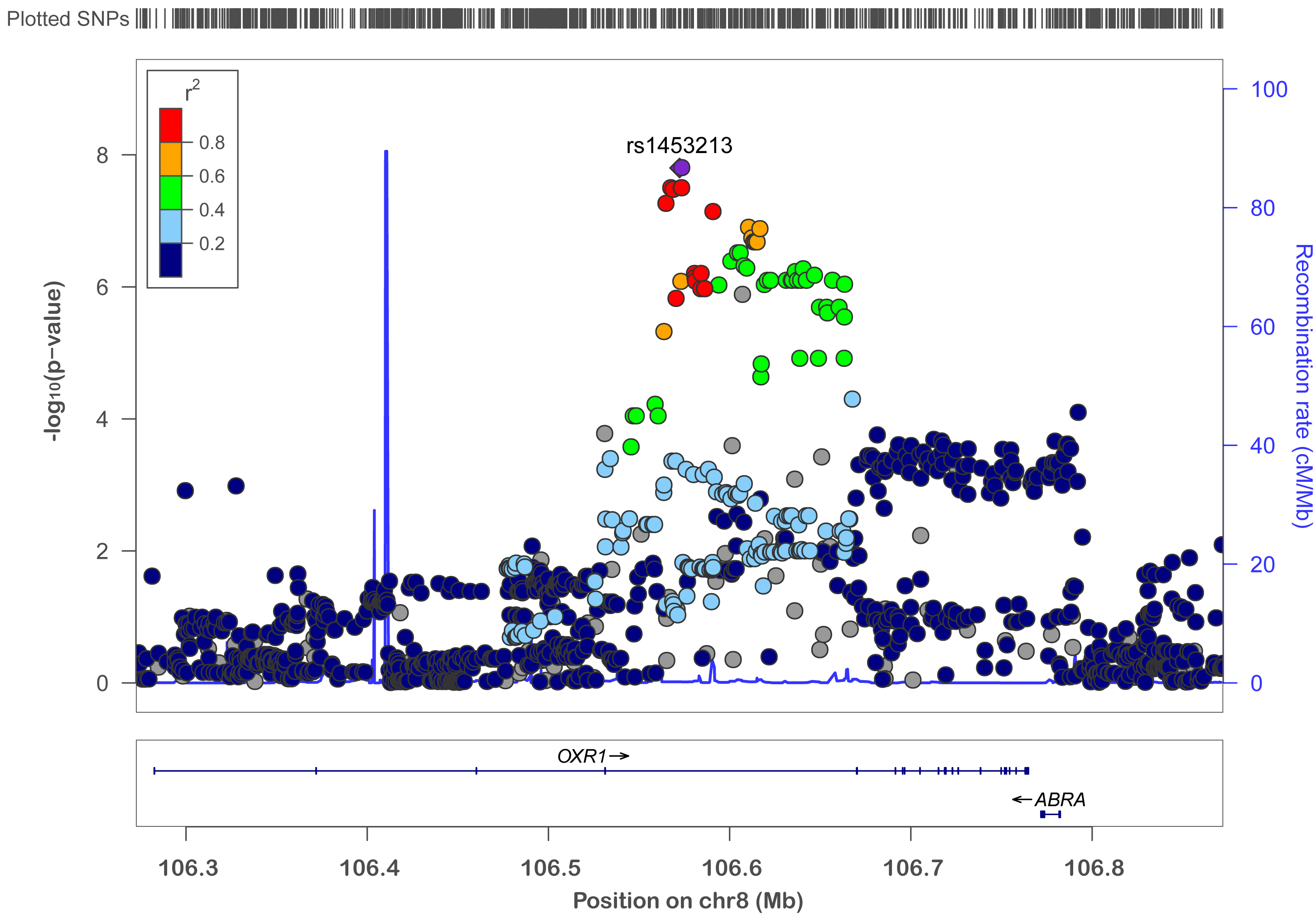

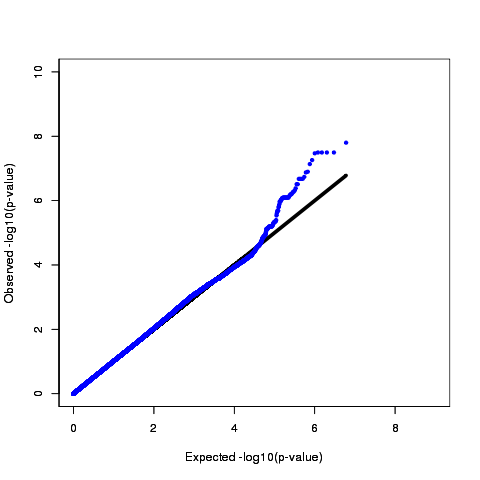

**Figure S4. Regional and Quantile-quantile (QQ) plots showing the associations between two top loci and taxa.** (A) Regional plot of *LOC150935* associated with phylum Actinobacteria, the index SNP rs62183161 (P=1.68 × 10^−9^) was showed in purple rhombus. (B) QQ plot with observed –log10 (p values) and the expected –log10 (p values) for phylum Actinobacteria. The genomic inflation factor is 1.019 (λ=1.019). (C) Regional plot of *OXR1* associated with family Prevotellaceae, the index SNP rs1453123 (P= 1.58 × 10^−8^) was showed in purple rhombus. (D) QQ plot with observed –log10 (p values) and the expected –log10 (p values) for family Prevotellaceae. The genomic inflation factor is 1.018 (λ=1.018).

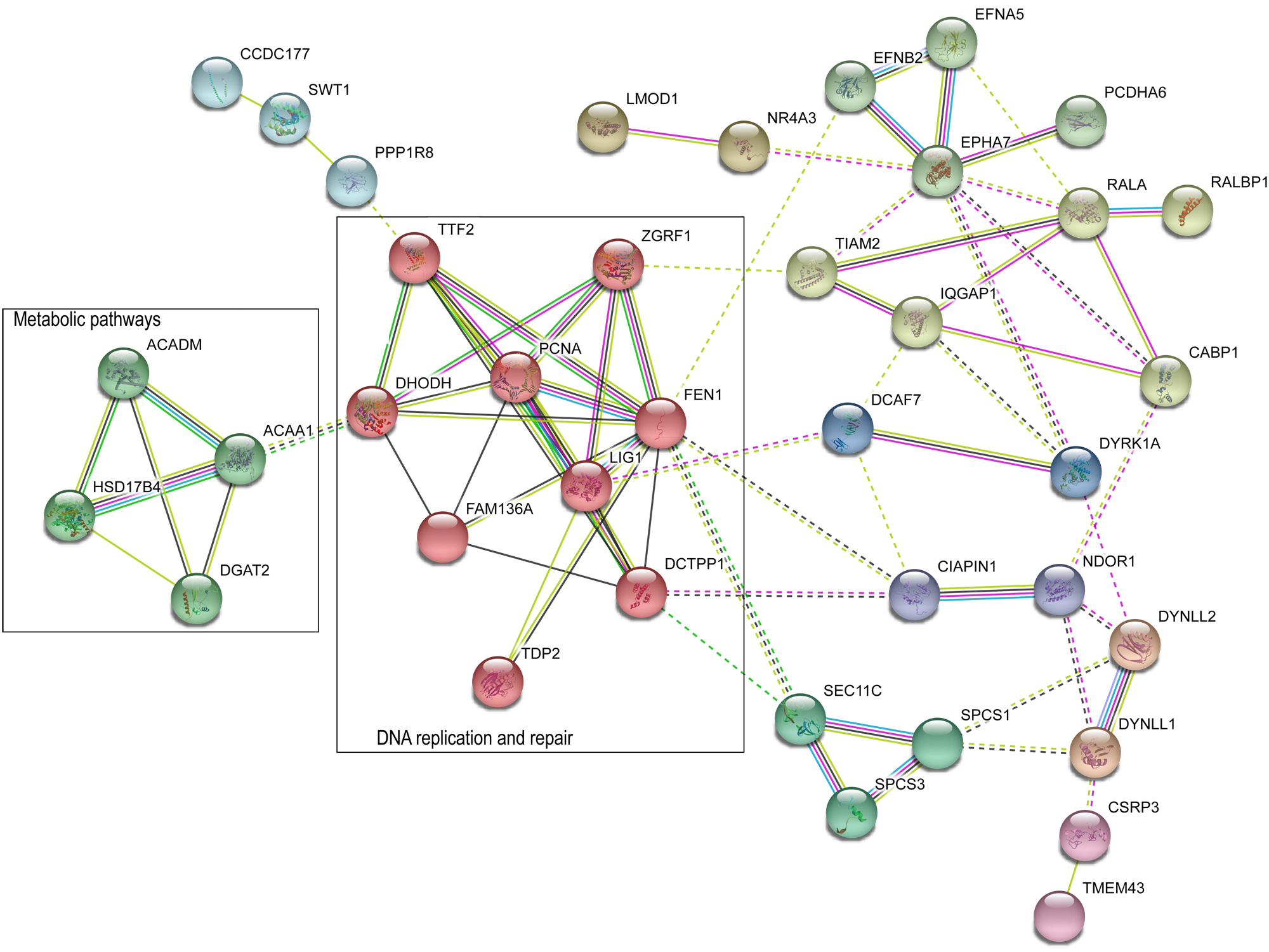

**Figure S5.** **Protein-protein interaction analyses.** The network contains 34 proteins and their 5 connected protein. Stronger associations are represented by thicker lines. Enrichment p-value=0.037. Two main pathways were marked with black rectangles, including metabolic pathways and DNA replication and repair pathways.

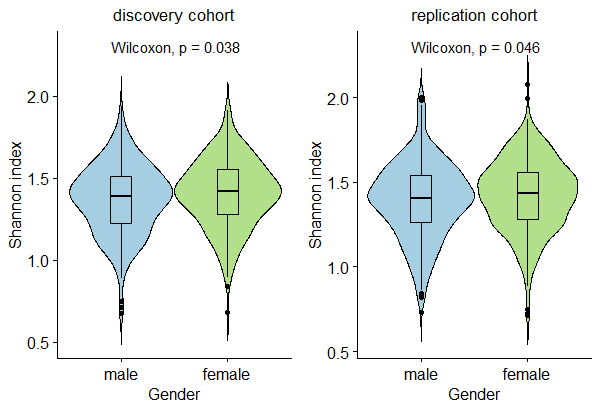

**B**

**A**

**Figure S6. Violin plots of sex differences on alpha diversity.** Alpha diversity was calculated for Shannon index based on genus-level relative abundance of taxa. Pairwise comparisons were performed using non-parametric test (Wilcoxon test). Both discovery and replication cohorts showed sex differences (Wilcoxon *P*=0.038 and *P*=0.046, respectively).
